# Impacts of stress and aging on spore health in *Schizosaccharomyces pombe*

**DOI:** 10.1101/2025.05.29.656811

**Authors:** Nicole L. Nuckolls, Michael T. Eickbush, Jeffrey J. Lange, Christopher J. Wood, Stephanie H. Nowotarski, Sarah E. Zanders

## Abstract

Most fungi can produce dormant, long-lived cells known as spores. Spores play a critical role in fungal biology and human health, but much about spores is unknown. Here, we investigate factors affecting spore fitness using the fission yeast *Schizosaccharomyces pombe* as a model. We found that storage conditions affect spore longevity, and that spore health declines over time. We identified a delay in dormancy breaking (germination), decreased asymmetry during cell division, and reduced stress tolerance as aging phenotypes. These results support that *S. pombe* spores are affected by both time and experiences during dormancy, highlighting critical features of spore biology and revealing parallels between aging in spores and aging in animal cells.

**Importance:** Fungi are integral to human health and well-being. Fungi also offer scientists easy experimental systems to study facets of life shared with other eukaryotes, including humans. Most studies of fungal cell biology focus on actively growing cells, but fungi can also produce dormant cells known as spores. Spores promote fungal survival and dispersal and are often the agents of infection in pathogenic fungi. In this work, we characterize the spores produced by an easy to study model fungus, *Schizosaccharomyces pombe*. We find that the longevity and the health of *S. pombe* spores declines over time and in response to heat stress. We characterize several traits associated with stressed and aged spores and identify parallels to aging cells in animals. This study expands the foundation for using *S. pombe* spores as a model system for fungal spore biology and as a model for aging of non-dividing cells.

## Introduction

Spores are specialized, dormant cells produced by most fungal species. Some fungi produce spores asexually from mitotically growing cells, some make spores sexually from the products of meiosis, and some can make both types of spores (*Wyatt et al., 2013, Dijksterhuis et al., 2019)*. While spores are diverse in form and DNA content, they are generally long-lived and tend to show enhanced resistance to stressors including enzymatic digestion, chemicals, heat, ultraviolet light, and desiccation (*Dijksterhuis et al., 2019; Ohtsuka et al., 2022; Ortiz & Hull, 2024).* Spores have a specialized, thick spore wall, as well as a dense cytoplasm enriched with protective molecules (e.g glycerol, mannitol, or trehalose) that give them enhanced stress resistance (*Ohtsuka et al., 2022; Plante et al., 2023; Sakai et al., 2024; Huang and Hull, 2017; Ortiz and Hull, 2024*). The viscous cytoplasm hinders the movement of molecules within the cell, reducing metabolic rate. Spores can further downregulate their metabolism via modulation of gene expression and through the polymerization of metabolic enzymes into aggregates (*Plante et al., 2023; Hugener et al., 2024)*.

Under permissive conditions, spores can undergo germination – the process of dormancy breaking. Germination ultimately results in the loss of the spore’s stress resistant features and the resumption of growth and cell division, although some stress resistance can persist for a few cell divisions in *S. cerevisiae* (*Gutierrez et al., 2018*). The exact stimuli that drive germination vary between fungi, but a carbon source (e.g. glucose) is sometimes required *(Ohtsuka et al., 2022; Hatanka and Shimoda, 2001; Sturm & Dworkin, 2015; Plante and Landry, 2021)*.

Germination of fungal spores involves isotropic swelling, outgrowth, and cell division and the process can be influenced by spore age, density, and environmental conditions (e.g. temperature, pH, nutrients) (*Ohtsuka et al., 2022; Ortiz and Hull, 2024; Veerana et. al.,2019; Punt et al., 2020; Krijgsheld et al., 2013; Bonazzi et al., 2014, Plante & Labbé, 2019; Plante et al., 2017*).

Spores play critical roles in fungal biology including facilitating dispersal and allowing fungi to survive environmental stresses (*Dijksterhuis et al., 2019*; *Ohtsuka et al., 2022; Ortiz and Hull, 2024).* Spores are extremely common in nature, and many pathogenic fungi, including *Cryptococcus* and *Aspergillus* species, generally infect humans via inhalation of spores (*Osherov 2012; Kwon-Chung and Sugui 2013; Wyatt et al., 2013; Dijksterhuis et al., 2019; McCartney and West, 2007; Ortiz and Hull, 2024).* In addition, plant pathogenic fungi (e.g. the basidiomycete plant pathogen, *Ustilago maydis*) can also spread via spores, resulting in crop losses and food spoilage (*Saville et al., 2012; Ortiz and Hull, 2024*). Understanding spore biology is therefore critical to human health and wellness.

More broadly, fungal spores offer an underexplored opportunity to understand dormancy – one of the most wide-spread cellular survival strategies *(Gremer and Sala, 2013; Miller et al., 2021)*. For example, plants, nematodes, rotifers, and tardigrades all have the potential to experience dormant life stages that contribute to long-term survival under changing, sometimes hostile, conditions (*Guidetti, Altiero, and Rebecchi, 2011; García-Roger et al., 2019; Vlaar et al., 2021; Penfield & MacGregor, 2017).* While humans do not have a systemic dormant state, many human cells are non-proliferative and can even enter a dormant state. These cellular transitions have critical health implications. For example, cancer cells can exhibit a treatment-resistant state of dormancy, where they become quiescent only to resume proliferation after treatment has eliminated primary tumors *(Phan & Croucher, 2020, Miller et al., 2021)*. However, our understanding of molecular mechanisms of cellular dormancy, in general, is limited.

Spores produced by the fission yeast *Schizosaccharomyces pombe* offer a highly tractable model system to explore spore biology and cellular dormancy (*Sakai et al., 2025*). *S. pombe* generally grows as haploid yeast cells. When *S. pombe* haploids are starved for nutrients, particularly nitrogen, two cells with opposite mating types can mate to form a diploid cell. Under persistent starvation, that diploid cell undergoes meiosis to generate four haploid cells that develop into spores (*Ohtsuka et al., 2022; Shimoda and Nakamura, 2004; Yamamoto, 1996*). *S. pombe* researchers have long known that *S. pombe* spores are generally viable for long periods when stored in water at four degrees Celsius. Still, the exact limits on longevity of *S. pombe* spores and the environmental factors affecting it are largely undocumented.

In this work, we use *S. pombe* spores to investigate factors affecting spore health and longevity. We found spore health is affected by storage temperature, highlighting that experiences during dormancy can impact future health. In addition, *S. pombe* spores display aging phenotypes, making them a tractable model for studying aging of non-dividing cells. Overall, our results expand our understanding of stress and aging during cellular dormancy.

## Results

### Storage temperature affects spore longevity

We used sexually produced spores from a homothallic (self-mating) strain derived from the commonly used *Schizosaccharomyces pombe* (Lindner isolate, *Hoffman et al., 2015*). We produced spores by plating haploid cells on malt extract agar (MEA, see methods) where the cells mate to form diploids, undergo meiosis, package the four meiotic products into spores, and release the spores from the sacks (asci) that hold them (Figure S1A). We isolated the spores using a standard protocol that involves treatment with ethanol and glusulase (a snail gut enzyme) to kill vegetative cells (see methods; *Smith, 2009*). Unless otherwise noted, we stored the spores in water and germinated them on solid, rich media (YEAS, see methods) at 32°C.

To understand how spore fitness might change over time, we first assayed spore viability at different storage temperatures: 4°C, 25°C, 32°C, and 37°C. In all conditions, we saw a decrease in spore viability over time, as measured by colony forming units (CFUs). The viability loss was slowed by storage in colder temperatures (Figures 1A and S1B-D). For example, spores stored at 37**°**C generally dropped under 50% viability by 10 days, while it took at least 80 days to drop to under 50% viability at 4**°**C (Figures 1A and S1B-D). This longevity at 4**°**C in water is less than what has been previously reported for *S. pombe* (35% viability after ∼1700 days; *Ohtsuka et al., 2022*), perhaps due to differences in strain background, reagents, or spore isolation protocols. Our observations of *S. pombe* spore longevity at 25°C (∼20% viability after 60 days) are, however, reasonably similar to previous reports for spores stored at 30°C (∼10% viability after 60 days; *Ohtsuka et al., 2022*). In addition, our longevity assays were reproducible across replicates, with some batch effects (Figures 1A and S1C; Table S1). Altogether, our results support the conclusion that temperature experienced during dormancy affects *S. pombe* spore survival. These results are also consistent with previous work showing that some spores from *Aspergillus, Neosartorya,* and *Penicillium* species lost viability at higher temperatures *(Gougouli and Koutsoumanis, 2013; Samapundo et al., 2007; Dagnas et al., 2017; Araujo & Rodrigues, 2004; van den Brule et al., 2022)*.

**Figure 1.**
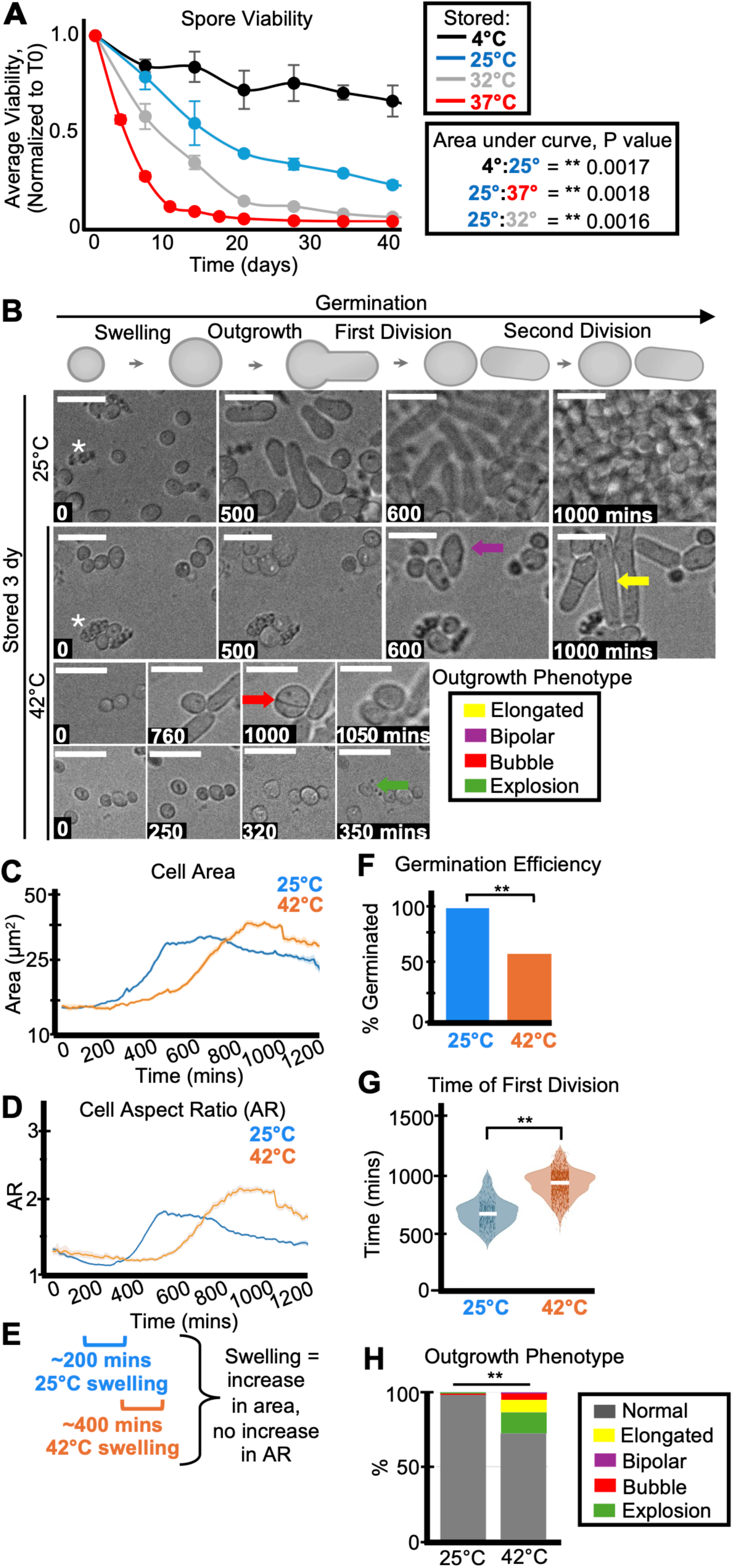
Storage temperature affects spore health and longevity. (A) One population of spores in water was split and stored at 4°C, 25°C, 32°C, or 37°C. The stored spores were then sampled over time to assay the number of colony-forming units (CFUs) on rich media (YEAS), which were normalized to the time zero starting point. The average of 3 replicates is shown and the error bars represent standard deviations. ** indicates p value < 0.01, comparing areas under the curves via t-test. (B) Timelapse microscopy images of spores that were stored 3 days at 25°C (top) or 42°C (bottom) prior to being plated on YEAS to induce germination at 32°C. The spores were imaged every 10 minutes for 1,200 minutes. Scale bars represent 10 μm. The colored arrows represent the following aberrant spore outgrowth phenotypes: elongated outgrowth (yellow), bipolar outgrowth (purple), bubble outgrowth (red), spore explosion (green). The white asterisks highlight debris. (C) Cell area (µm^2^) and (D) cell aspect ratio (AR) during germination of spores stored for three days at 25°C (blue) or 42°C (orange). The shaded area around each line represents standard error. In some instances, the standard error is thinner than the line. Note that these cell areas provided by the deep learning algorithm are slightly larger than the actual cell areas (see methods). (E) Swelling is defined as an increase in cell area without an increase in aspect ratio (departure from circularity). The brackets indicate approximate windows of swelling derived from the plots in C and D. (F) The fraction of spores manually observed to germinate by the end of the 1,200-minute timelapse. N>50 spores per sample. ** indicates p-value < 0.01 via Fisher’s exact test. (G) Time of first division (time when the AR of a cell exceeded 3) of spores after storage for 3 days at 25°C or 42°C. ** indicates p- value <0.01 via ANOVA test. N>100 spores per sample. (H) The percentage of spores displaying the indicated outgrowth phenotypes. N>50 outgrowths per sample. ** indicates p- value < 0.01 via Fisher’s exact test.

## Storage temperature affects spore health

To assay how the quality of spores, beyond their viability, might change over time at different storage temperatures, we observed cytological features of germination in time-lapse images. We plated spores on solid, rich media and imaged them every 10 minutes for at least 1,000 minutes (∼20 hours) at 32°C (Figures 1B and S1E). We classified spores as successfully germinated if they completed one cell division within this timeframe. To facilitate scoring the timelapses, we trained a deep learned algorithm to identify spores. We then used standard image processing approaches to measure the area and aspect ratio (ratio of the long to short axis of an ellipse fit to the object) of each tracked cell in each time point. The first division (or successful germination) was defined as the time when a cell’s aspect ratio exceeds 3 (see methods). We also scored at least 30 cells per sample manually to confirm the algorithm outputs. We noted that the fraction of spores we observe to germinate is often higher than expected based on our colony forming unit (CFU) measurements. This is likely because spores can lose integrity upon death and are not recognized and assayed in the cytological experiments. Consistent with this, we see debris accumulate in the spore storage tubes (as a layer above the spore pellet) as well as in the background of our timelapses of samples where CFUs have dropped (asterisks in Figure 1B). In addition, some spores may be capable of completing one cell division but may not be capable of generating a colony. Because of these factors, we consider the colony forming unit assays as a better measurement of viability, while the cytological analyses reveal additional phenotypes.

At the standard storage temperature of 4°C, we found that fewer of the visible spores germinated in older compared to younger samples, as expected from our CFU analyses (Figures S2A-D, S2F-I and S3A-D). Despite the loss in viability, we found that the viable spores largely remained healthy during storage 4°C, with no gross defects in germination morphology. In two experiments, we found that older spores (65 and ∼400 days) were slower to swell (as defined by an increase in spore area without an increase in aspect ratio) and germinate than 1- day-old samples (Figures S2A-C, E and S3A-C, E). We did not, however, observe a difference in swelling or germination timing of spores from a third experiment that compared spores stored for 15 and 397 days at 4°C, suggesting that changes observed at 4°C in germination timing are small and thus within the range of experimental variability of these phenotypes (Figures S2F-H, J).

We next assayed the health of spores stored at warmer conditions, first comparing 25°C (mild conditions) to 42°C (heat stress) for 3-days. As expected, we found that fewer of the spores stored at 42°C germinated relative to the spores stored at 25°C (Figure 1F). The spores stored at 25°C were also faster to swell and divide than spores stored at 42°C (Figures 1C-E, G and S4A-B). We observed delayed germination of spores stored at 42°C across multiple timelapses, new spore isolations, and distinct germination conditions (Figure S5). These data show that 42°C-storage is reproducibly detrimental to spore health. Previous work found that larger spores germinate faster (*Padilla et al., 1975*), so we tested if the germination delay in the spores stored at 42°C was associated with decreased size. This was not the case, as the spores stored at 42°C were either not significantly different in size, or slightly larger than those stored at 25°C (Figures S4C and S5G).

We noticed that spores stored for three days at 42°C often displayed abnormal outgrowth phenotypes (Figure 1B arrows, quantified in Figures 1H and S5E). Generally, germinating spore outgrowths are ∼4.5 µm prior to the first division, but some spores stored at 42°C produced longer outgrowths (as long as 20 µm), suggesting cell cycle defects (yellow arrows, Figure 1B; *Wood and Nurse, 2015; Nurse et al., 1976*). In addition, we observed an increase in polarity defects amongst the spores stored at 42°C for three days (*Bonazzi et al., 2014; Wei et al., 2023*). Specifically, we observed “bubble” like divisions, where the cells remain circular and divide in the middle of the “bubble” (red arrow, Figure 1B). We also observed increased bipolar outgrowths from spores stored at 42°C, where the spore grows from both ends, rather than asymmetrically from one side (purple arrow, Figure 1B). Finally, we observed an increase in “exploding” spores (green arrow, Figure 1B) that disintegrate during germination amongst those stored at 42°C.

We also assayed spores stored at a milder heat stress, 37°C, for 15-28 days. Like the spores stored at 42°C, we saw reductions in germination efficiency of the visible spores, delayed germination, and an increase in abnormal outgrowth phenotypes in the spores stored at 37°C, relative to those stored at 25°C. These effects were observed with spores germinated on rich media at both 32°C and 37°C and on synthetic media at 32°C (Figures S6-S8).

To test if storage under heat stress affected spore morphology, we imaged spores using Scanning Electron Microscopy (SEM) and Scanning Transmission Electron Microscopy (STEM). We saw no differences in the external appearance of spores stored for 3 days at 25°C or 42°C (Figure S9A-B). We did see, however, that more of the spores stored at 42°C, relative to those stored at 25°C, had lost the dark-straining (i.e. electron dense) appearance of the cytoplasm (Figure S9C-D). All together, these observations demonstrate that storage in heat-stressed conditions negatively impacts spore health.

## Storage medium affects spore longevity

Spores isolated from *Saccharomyces* and *Aspergillus* can incorporate labeled nucleotides and amino acids, suggesting spores may carry out some transcription and translation (*Maire et al., 2020; Wang et al., 2021).* We, therefore, wanted to test if a nutrient-dense storage medium affects *S. pombe* spore health by comparing spores stored in water to those stored in a solution of 0.5% yeast extract (YE). We stored the spores at 4°C, 25°C, and 37°C and measured spore viability over time by assaying colony forming units. Spores stored in YE at 25°C and 37°C generally survived longer than matched samples stored in water (Figure 2A-B). At 4°C, we observed slightly higher viability of spores stored in YE after ∼300 days in one experiment (Figure S10A), but the replicate experiment was terminated after 109 days and there was no viability difference between spores stored in YE and those stored in water (Figure S10B).

**Figure 2.**
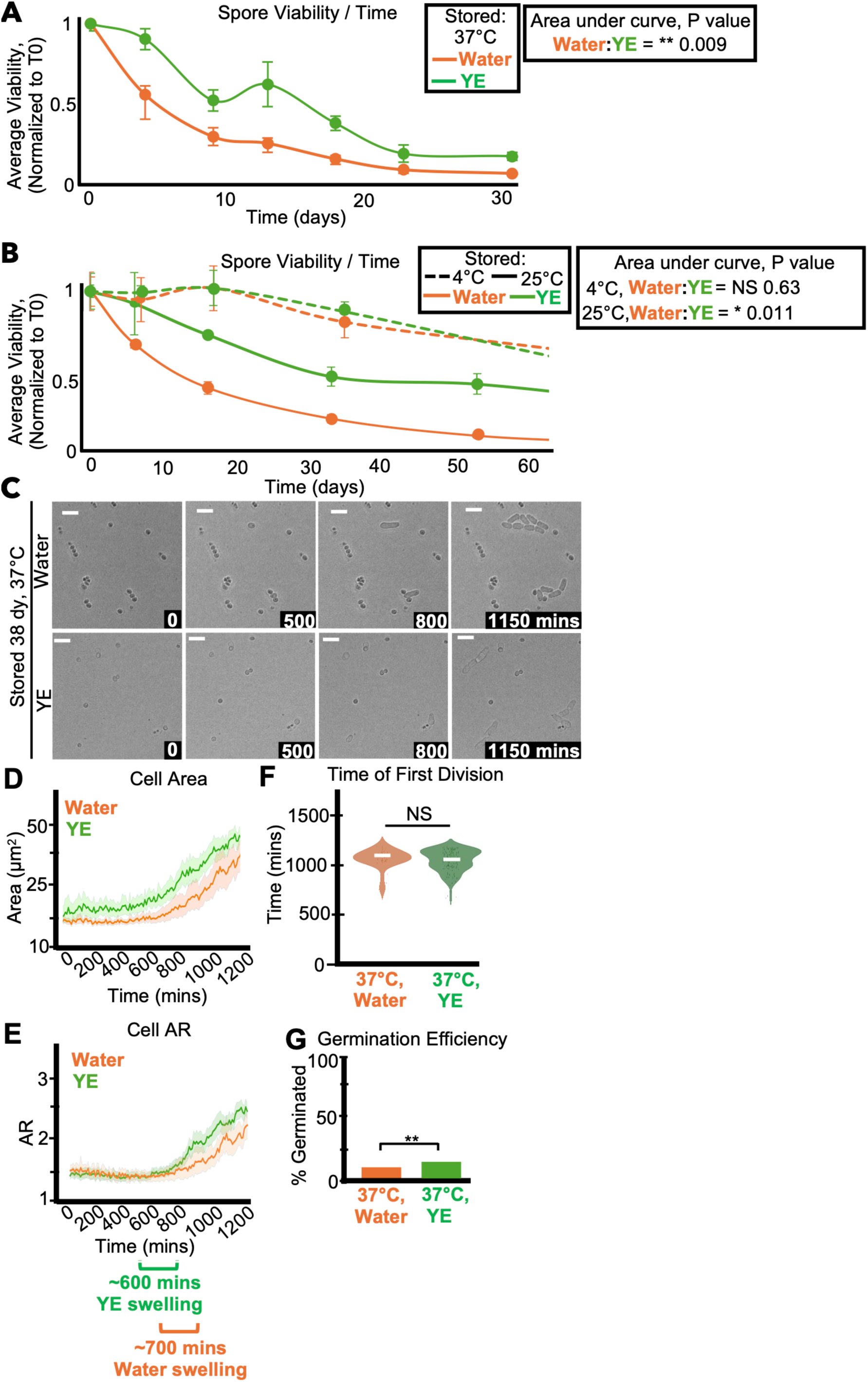
Storage media affects spore longevity. (A) One population of spores was split and resuspended in either 0.5% yeast extract solution without glucose (YE) or water and then stored at 37°C. The same experiment was performed in (B), but the spores were stored at 4°C or 25°C. In both experiments, the stored spores were then sampled over time to assay the number of colony-forming units (CFUs) on rich media (YEAS), which were normalized to time zero starting point. We show the average of 3 replicates and error bars represent standard deviation. ** indicates p value < 0.01, * indicates p value < 0.05, comparing areas under the curves via t- test. (C-G) Imaging analyses of spores stored for 38 days at 37°C in either water or yeast extract solution before being plated on rich media (YEAS) to induce germination at 32°C. Cells were imaged every 10 minutes for 1,200 minutes. (C) Timelapse microscopy images of germinating spores. The time in minutes is displayed and the scale bars represent 10 μm. (D) Cell area (µm^2^) and (E) cell aspect ratio (AR) during germination. The shaded area around each line represents standard error. In some instances, the standard error is thinner than the line. Note that these cell areas provided by the deep learning algorithm are slightly larger than the actual cell areas (see methods). (F) Time of first division (time when the AR of a cell exceeded 3). ** indicates p-value <0.01 via ANOVA test. N>100 spores per sample. (G) The fraction of spores manually observed to germinate by the end of the 1,200-minute timelapse. N>50 spores per sample. ** indicates p-value < 0.01 via Fisher’s exact test.

Importantly, we did not identify what feature(s) of YE supports spore longevity better than water at warmer temperatures. We speculate it is the presence of nutrients, but we did not rule out any potential contributions of osmolarity or slight pH differences.

We also assayed the effects of storage media on spore health by cytologically comparing cells stored at 4°C, 37°C, and 42°C in either YE or water. We observed a trend of more and faster germination amongst spores stored in YE relative to those stored in water. However, these effects (except germination efficiency after 38 days at 37°C) were not statistically significant in individual experiments (Figures 2C-2G and S10-S12). Together, our results suggest that storage in YE significantly affects spore survival at warmer temperatures, but does not significantly influence the health of the viable spore population.

## Age affects spore health

Our data (Figure 1A) demonstrates that spores lose viability over time and our cytological observations of spores stored at 4°C suggest that spore health can decline over time, but the effects were small and varied between experiments (Figures S2 and S3). We predicted any age-associated effects would manifest more quickly and be more obvious with spores stored at warmer temperatures. We therefore compared germination phenotypes of spores stored at 25°C for 7 days, 116 days, or 176 days. As expected, significantly fewer of the visible 116- and 176-day-old spores germinated than the 7-day-old spores (Figure 3A, D). The aged spores were also significantly slower to swell, outgrow, and divide than the 7-day-old spores (Figure 3B-C, E). The differences in germination time were repeatable across distinct spore isolations (Figure S13), not due to size differences between spores (Figures 3G) or spore concentration (Figure S14) and were observed across different germination conditions (Figures S15 and S16). We also noted significantly more abnormal germination morphologies in the aged spores (Figures 3F and S13F), like those observed in the young spores stored at 3 days at 42°C (Figure 1H and S5E).

**Figure 3.**
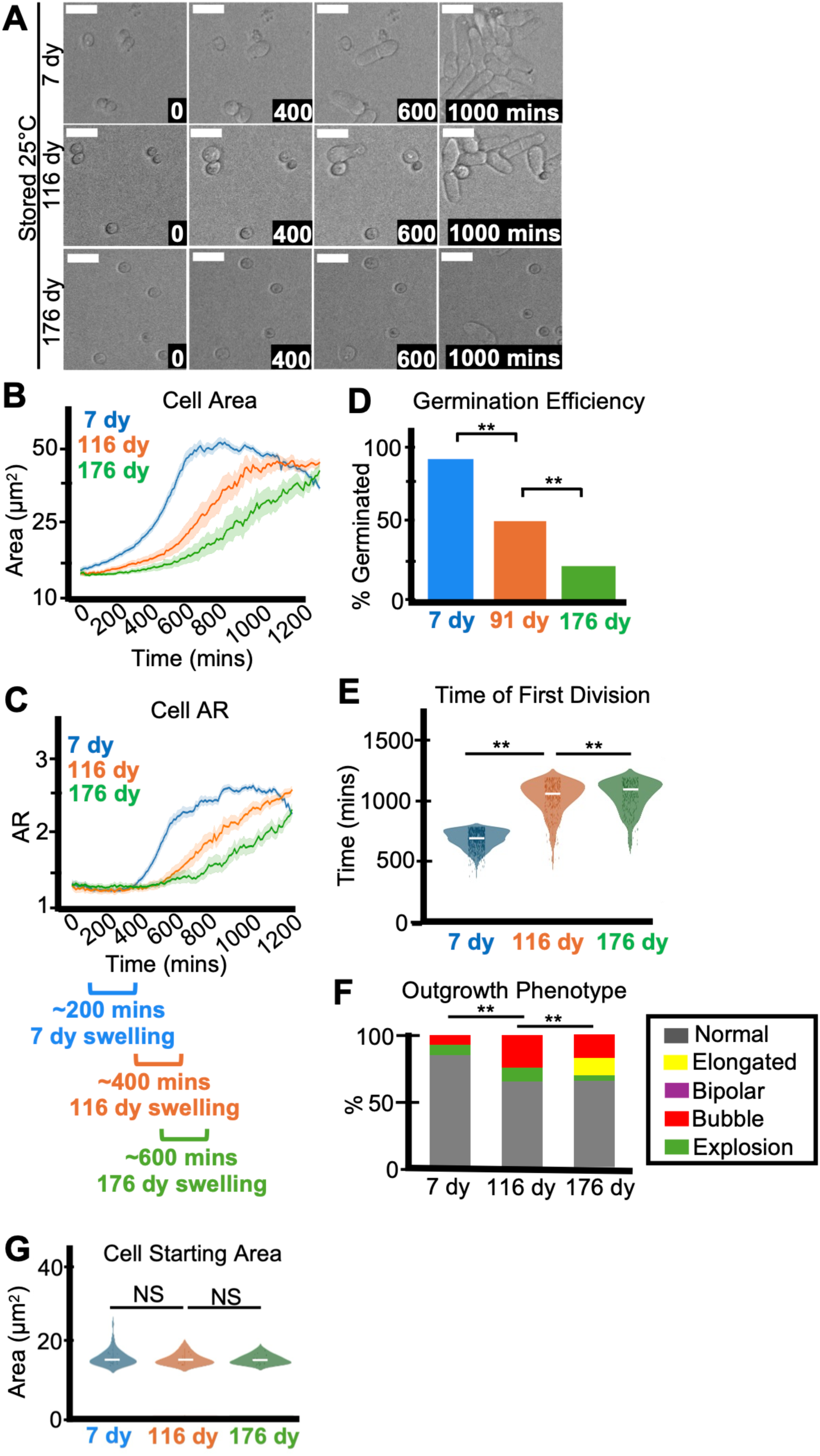
Age affects spore health. Imaging analysis of spores stored at 25°C for 7, 116, or 176 days in water before being plated on rich media (YEAS) to induce germination at 32°C. Cells were imaged every 10 minutes for 1,200 minutes. (A) Timelapse microscopy images of germinating spores. The time in minutes is displayed and scale bars represent 10 μm. (B) Area (µm^2^) and (C) aspect ratio (AR) during germination. The shaded area around each line represents standard error. In some instances, the standard error is thinner than the line. Note that these cell areas provided by the deep learning algorithm are slightly larger than the actual cell areas (see methods). (D) The fraction of spores manually observed to germinate by the end of the 1,200-minute timelapse. N>50 spores per sample. ** indicates p-value < 0.01 via Fisher’s exact test. (E) Time of first division (time when the AR of a cell exceeded 3). ** indicates p-value < 0.01 via ANOVA test. N>100 spores per sample. (F) The percentage of spores displaying the indicated outgrowth phenotypes. N>50 outgrowths per sample. ** indicates p-value < 0.01 via Fisher’s exact test. (G) Starting area (µm^2^) of spores. N>50 spores per sample. NS indicates not significant via ANOVA test.

Finally, similar to heat stressed spores, we observed significantly more spores with light-staining (i.e. less electron dense) cytoplasm in older spores compared to young (Figure S9C-D). We also saw more deflated, shrunken spores in the older samples than the younger (Figure S9A-B). In all, these observations demonstrate that age can negatively impact spore health.

## Age and stress affect asymmetry of spore germination

Previous work demonstrated intracellular asymmetry in germinating *S. pombe* spores in that vacuoles are preferentially retained in the round end of germinating spores, as opposed to the outgrowth (*Plante et al., 2017)*. We wondered if the decreased asymmetry we observed in the shape of germinating heat stressed (Figures 1B, 1H, S5E, S6E, S7E, and S8E) and aged spores (Figures 3F and S13F) extended inside of the cells to the vacuole. To test this, we assayed the partitioning of dye-stained (FM4-64) vacuoles during germination of spores stored at 25**°**C for 16 days, 105, and 140 days, as well as 37**°**C for 16 days (Figure 4A). We saw that the 25**°**C-stored, 16-day-old and 105-day-old spores retained most (64 and 60%, respectively) of the vacuole signal in the spore, with less FM4-64 signal in the outgrowth (Figure 4B and 4C). However, the 25**°**C-stored 140-day-old and 37**°**C-stored, 16-day-old spores had significantly more vacuole straining entering the outgrowth (Figures 4B-C). This phenotype of decreased vacuole asymmetry at the first division in older and heat stressed spores persisted between spore preparations (Figures S17 and S18) indicating that spores’ ability to asymmetrically segregate vacuoles can decline due to heat stress and due to aging.

**Figure 4.**
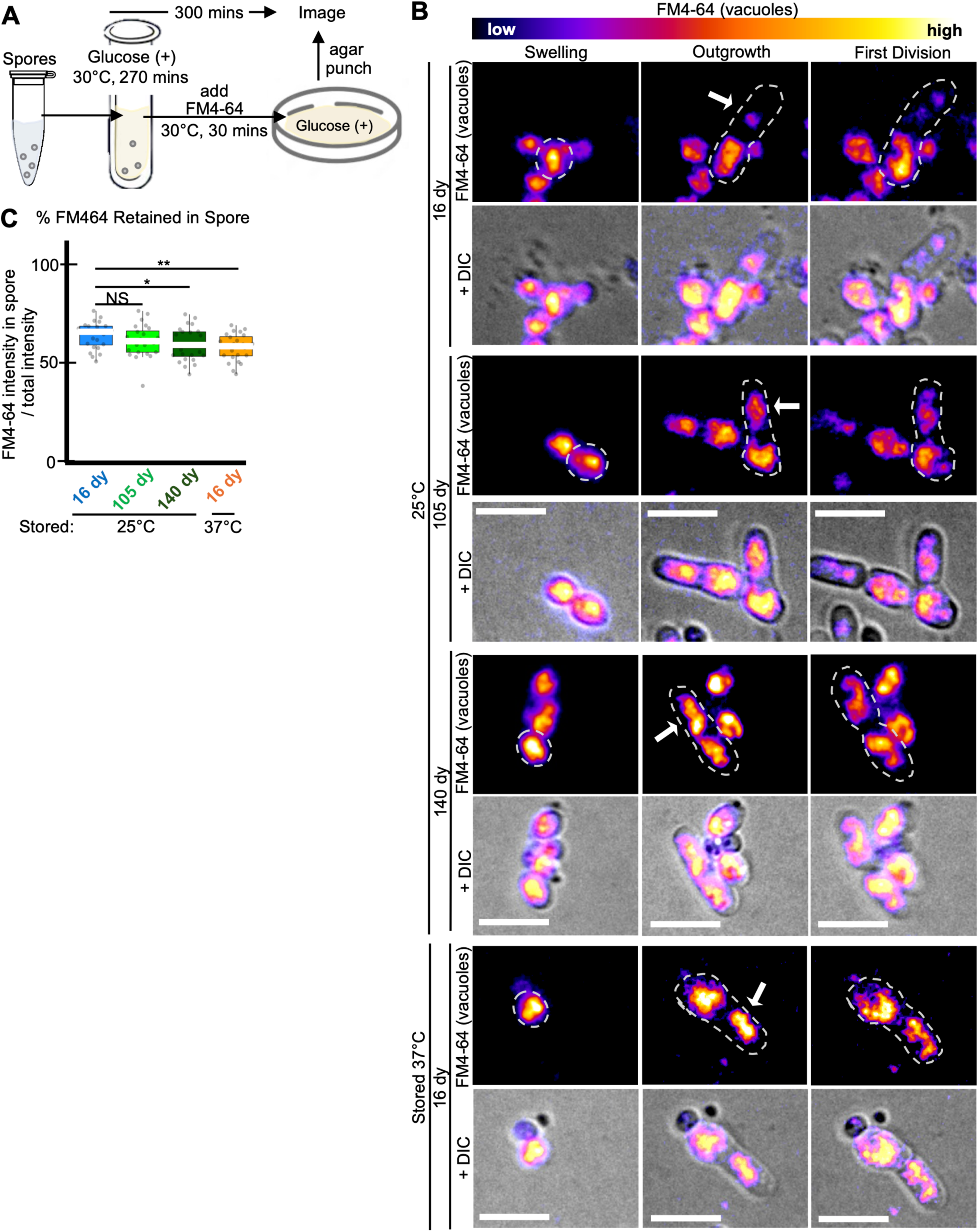
Age affects asymmetry of vacuole segregation during germination. Imaging analyses of vacuole segregation during germination of spores previously stored for 16, 105, or 140 days at 25°C in water or for 16 days at 37°C in water. (A) Cartoon describing experimental approach. Spores were resuspended in rich liquid media (YES) at 30°C for 270 minutes to induce germination. The vacuole dye FM4-64 was then added, and the cells were grown for an additional 30 minutes before being plated on solid rich media (YEAS). The cells were then imaged on the agar at 32°C using timelapse microscopy. (B) Timelapse microscopy images of FM4-64-stained spores for 20 hours. The brightness and contrast are not the same for all images but were adjusted so the spore bodies appeared to have similar levels of signal. Scale bars represent 10 μm. (C) Percentage of the total FM4-64 signal retained in the spore-body verses the germ tube outgrowth. N> 30 spores. ** indicates p-value <0.01, * p-value <0.05, NS indicates not significant, via Fisher’s exact test. Error bars show standard error.

## Prior stress and aging affect spore stress tolerance

A key spore phenotype is that they display resistance to stresses (*Arellano et al., 2000; Coluccio et al., 2008; Egel, 1977; Fukunishi et al., 2014; Shimoda, 1980*). We hypothesized that past experiences and age could affect stress resistance in spores. To test this, we counted the number of colony-forming units produced by spore populations before and after heat stress (55°C) and before and after exposure to Ultra-Violet (UV) light (see methods). We found that a significantly larger fraction of the spores stored at 25°C survived both heat and UV shock compared to age-matched spores stored at 37°C (Figures 5 and S19). Similarly, we found that young spores had higher resistance to heat and UV stress than older spores (Figures 5 and S19). These experiments demonstrate that age and storage conditions influence a spore’s ability to tolerate stress.

**Figure 5.**
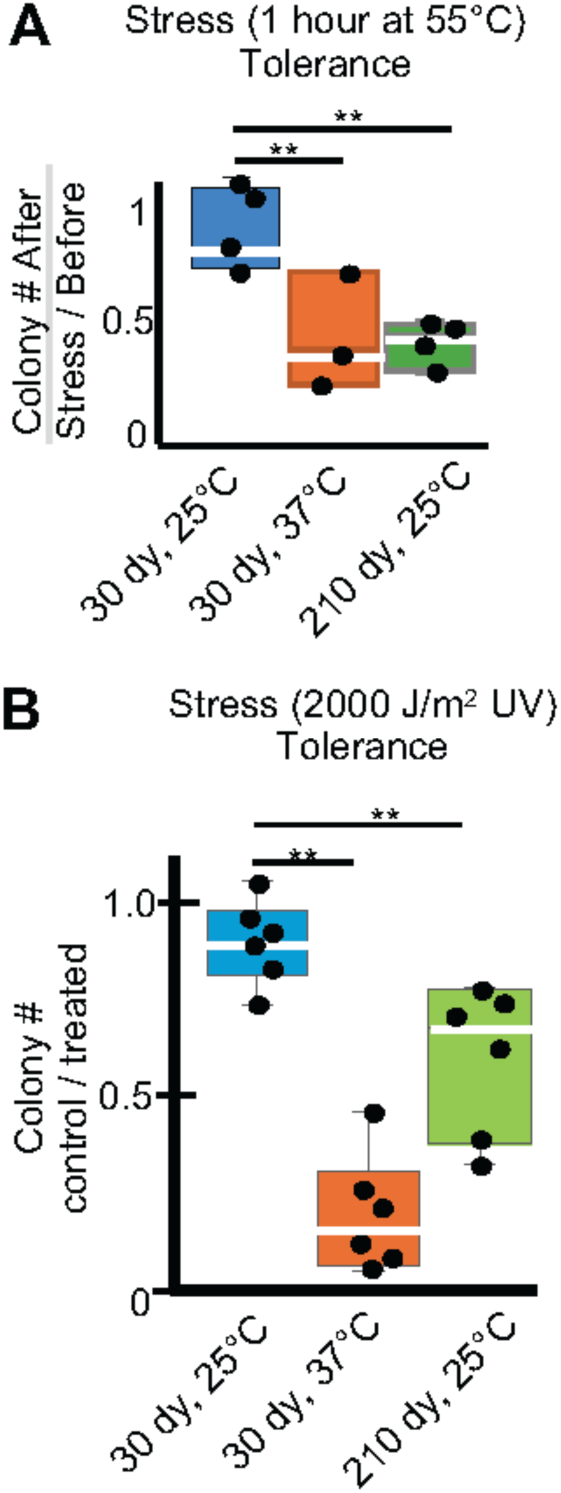
Age and stress affect spore stress tolerance. The colony forming units (CFUs) of spore samples previously stored for 30 days at 25°C or 37°C or for 210 days at 25°C were determined by plating on rich media (YEAS) at 32°C both before and after an acute stress. (A) The ratio of surviving CFUs before and after a 1-hour heat shock of 55°C or (B) Exposure to 2000 μJules UV. ** indicates p-value < 0.01 via t-test. N≥3 replicates.

## Discussion

Understanding spore formation, maintenance and germination is crucial for understanding the biology of fungi. But much remains to be learned about these questions, even in intensely studied and tractable model systems like *S. pombe* (*Sakai et al., 2025*). Here, we aimed to help fill that gap by investigating influences on health and longevity of *S. pombe* spores. We established parameters for *S. pombe* spore longevity under varying conditions (Figure 1A).

Similar to work in *Penicillium*, *Aspergillus*, and *Absidia*, we found that storage temperature significantly impacts health and longevity (*Gougouli and Koutsoumanis, 2013; Samapundo et al., 2007; Dagnas et al., 2017; Araujo & Rodrigues, 2004; Conner,Beauchat, and Chang 1987; Demming et al., 2011*).

To reduce the labor associated with studying germination, we improved our previously developed artificial intelligence-assisted image scoring approaches to automate scoring of the key germination phenotypes (swelling and first division timing, *Billmyre et al., 2022*). In addition, we identified several phenotypes associated with spore stress and aging: decreased germination efficiency, delayed germination, and altered spore outgrowths.

Consistent with standard practices within the *S. pombe* field, we found that some spores could maintain viability for long periods when stored in water at 4°C. We were surprised to find, however, that a sizeable fraction of spores stored at 4°C died within 100 days of storage under our laboratory conditions. Still, the spores that remained viable at 4°C appeared grossly normal and showed mild, or no obvious heath defects. Spores stored at higher temperatures, however, exhibited accelerated loss of health and viability.

For future analyses, we are particularly interested in the causes and consequences of the breakdown in vacuole retention in aged spores. We speculate that the asymmetry of the first division after germination may promote rejuvenation of the lineage founded by the spores. This idea is based on work in *Saccharomyces cerevisiae* (budding yeast), which has a similar asymmetric division style as the typical first and second division of *S. pombe* spores. In budding yeast, debris, (e.g. misfolded proteins/RNAs, extrachromosomal circles, and aggregates) are retained in the mother cell during cell division. This asymmetric cytoplasmic inheritance promotes the rejuvenation of the daughter cell (the bud) and this process is less efficient in aged mothers (*Smith et al., 2015; Lippuner et al., 2014; Steinkraus et al., 2008*). *S. pombe* spores could similarly use asymmetric division to purge debris in the first daughter cell. If so, this process is breaking down over storage time in spores.

Parallels between spore germination and animal stem cells are also notable. First, the asymmetric spore germination phenotype is reminiscent of the asymmetric divisions of human stem cells, where one daughter cell retains stem cell properties and the other differentiates into a specialized cell. Interestingly, asymmetric inheritance of lysosomes (the equivalent of fungal vacuoles) is an important aspect of this differentiation. However, older stem cells tend to lose their ability to divide asymmetrically and produce two daughter cells with similar characteristics, causing tissue decline associated with aging (*Yamashita et al., 2010; Mi et al., 2022*). In addition, we found that aged spores are slower to break dormancy than younger spores. This also mirrors an aging phenotype in human stem cells, where aged, non-dividing stem cells become more resistant to activation, generating a progressive age-associated delay in progressing through the cell cycle (*Viale et al., 2009; Sottocornola et al., 2012*). These similarities between spores and stem cells suggest that discoveries about experimentally tractable fungal spores may sometimes reflect deeply conserved biology and thus be broadly relevant.

We aspire that this study provides parameters and phenotypes that will be helpful in future studies addressing the molecular mechanisms underlying spore longevity and stress resistance phenotypes in the highly tractable *S. pombe* system. We anticipate at least some of these mechanisms may be shared with other fungal spores and other non-dividing eukaryotic cell types.

## Materials and Methods

### Inducing sporulation

We cultured SZY643 (*h90, leu1-32, ura4-D18; Nuckolls et al., 2017).* in 5 mL of YES rich media (0.5% yeast extract, 3% glucose, 250 mg/L of adenine, lysine, histidine, leucine, and uracil) overnight at 32°C with shaking. We then diluted the culture to 10^-3^, 10^-4^, and 10^-5^. We plated 100 µL of the 10^-3^ dilution on 5 separate Malt Extract Agar plates (MEA, 2% Difco Bacto Agar, 30 g/L Bacto-malt extract, 5 mg/L adenine, histidine, leucine, uracil pH adjusted to 5.5 with NaOH) to induce mating and sporulation. We also plated 100 µL of the 10^-4^ and 10^-5^ dilutions on rich media, YEAS (same as YES above with 2% agar). We counted the colony forming units on the YEAS plates after 3-5 days to estimate the number of cells plated on the MEA. We left the MEA plates at 25°C for 9 days. By day 9, the cells will have (1) self-mated, (2) undergone meiosis to produce spores, and (3) naturally broken down the spore-sac to release the spores (Figure 1A). Using a cell scraper (Fisher Scientific: 353086) and water, we harvested the total mass of all spots off the 5 MEA plates into a 50 mL conical tube in 20 mL water and continued to spore isolation methods (below).

### Isolation of spores

We killed vegetative cells to isolate spores using glusulase, followed by ethanol (*Smith, 2009)*. We added 100 µL of glusulase (Sigma-Aldrich: G0876) to the 20 mL of spores in water and placed the tubes at 32°C overnight with shaking. The next day, we added 20 mL of 60% ethanol for 10 minutes at room temperature. We spun to pellet, washed twice with 25 mL water, and resuspended the spores in 50 mL water. We then stored the spores in small aliquots in water (unless otherwise noted) at various temperatures. To limit phenotypes introduced by repeated vortexing and to prevent contamination, each aliquot was sampled once or for one day of experiments.

### Longevity Assays

We plated dilutions of spores onto YEAS counted the colonies after 3 and 5 days of growth 32°C (unless otherwise noted).

### Vacuole, FM4-64 staining

We found that dormant spores did not efficiently take up the dye FM4-64 (Invitrogen:T13320), but that the cells became permeable to the dye during germination. We therefore placed spores in YES liquid media for 270 minutes at 30°C with shaking to induce germination prior to adding FM4-64 (Invitrogen:T13320) into the YES media to a concentration of 10 µM. We then incubated the cells an additional 30 minutes shaking at 30°C to stain (Figure 5A). We then washed the germinating spores twice with water and proceeded to time-lapse microscopy of germination.

### Timelapse microscopy

The samples used in each experiment are described in Table S2. Some spore preparations were used in multiple experiments: e.g a distinct aliquot from a spore preparation used in one experiment as a young sample could later be used in a subsequent experiment as an aged sample. We diluted the spores to 10^-3^ in water. We then plated 100 µL of the 10^-3^ dilution onto germination media (media containing glucose, usually YEAS, with the exception of Figures S3, S5, S7, and S15 which utilize PMG [3 g/L potassium hydrogen phthalate, 2.2 g/L Na_2_HPO_4_, 3.75 g/lL glutamic acid, monosodium salt (Sigma G-5889), 20 g/L glucose, 20 mL/L salts (0.26 M MgCl_2_.6H_2_0; 4.99 mM CaCl_2_.2H_2_0, 0.67 M KCL, 14.1 mM Na_2_SO_4_) , 1 mL/L vitamins (1000x Stock: 2.20 mM pantothenic acid, 81.2 mM nicotinic acid, 55.5 mM inositol, 40.8 uM biotin), 0.1 mL/L minerals (10,000X Stock:80.9 mM boric acid; 23.7 mM MnSO_4_; 13.9 mM ZnSO_4_.7H_2_0; 7.40 mM FeCl_2_.6H_2_0; 2.47 mM molybdic acid; 6.02 mM KI; 1.60 mM CuSO_4_.5H_2_0; 47.6mM citric acid), and 2% agar]. We then took a 0.6 cm diameter punch of the agar and placed it upside down into an 8 well uncoated dish (Ibidi: 80826), trapping the cells between the agar and the dish for imaging. We then imaged the cells on a on a Ti2 widefield microscope (Nikon) through a 60x Plan Apo (NA 1.4, Nikon) oil objective. We imaged every 10 minutes for at least 1,000 minutes (∼20 hours), collecting transmitted light using the Prime 95B camera (Photometrics), using perfect focus on a single Z-slice. For the FM4-64 timelapse, we collected the FM4-64 signal (and transmitted light) every 15 minutes for 1,000 minutes (∼20 hours) on the same microscope and objective using an mCherry optical configuration with excitation centered at 586 nm and with the emission collected through an EM610/75 filter, again using perfect focus on a single Z-slice.

### Deep learning algorithm

We implemented a custom-trained Cellpose (version 3, (*Stringer et al., 2021*) model for spore tracking and measurements (area and aspect ratio). To create a training set, we labeled all cells in 35 random transmitted light images of spores to make image-mask pairs. Using the training set and starting from the pretrained cyto2 model, the Cellpose training module ran for 1000 epochs with an initial learning rate of 0.005 and a weight decay of 0.00001. We saved the model after training and used it for inference in further image analysis. The training and analysis scripts were custom written in python and the training computation was performed on an Ubuntu Linux 18.04 workstation with an NVIDIA RTX A6000 GPU. The scripts we generated are all publicly available in the Stowers Original Data Repository (ODR) at https://www.stowers.org/research/publications/LIBPB-2559.

### Plot generation

We used the area and aspect ratio (AR) measurements calculated by the algorithm for all data except the area of the area of germinated spores, which was measured manually (described below). If a timelapse temporarily lost focus, the data at that timepoint appears as a sharp dip or peak in the data (example: Figure S3, 438-day-old sample at 650 minutes). If a timelapse lost focus for more than 5 timepoints, we excluded the data from analysis. To plot time to germination, we scored only unique particles with an AR less than 1.4 in time point zero (these are circular and therefore likely spores). We then designated when that object reached an AR greater than 3.0 as the time of the first division. This cutoff was determined by manually comparing AR and division status of >50 cells, which tend to divide at a predictable size and had an average aspect ratio of 3.1 at division.

To plot germination efficiency, we scored using data collected manually by eye. We scored at least 100 spores per sample and counted how many successfully underwent first division before the end of the timelapse. We then divided that number by the total spores scored. We also scored the outgrowth phenotype manually by eye. We counted at least 50 outgrowths per sample and determined if it was normal, elongated (more than 6 μm), bubbled, exploded, or bipolar. If a spore was both bipolar and elongated, we counted it as bipolar.

To plot the starting area for all spores, we used data collected by the deep learning algorithm. We calculated the area (µm^2^) of unique particles with an AR less than 1.4 in time point zero. Note that the cell areas provided by the deep learning algorithm (e.g. starting spores ∼10-15 μm^2^) are larger than the actual cell areas (e.g. starting spores ∼7-9 μm^2^) due to the automatically-generated cell masks extending slightly beyond the cell boundaries. To determine the starting area for germinated spores, we scored using data collected manually. We calculated the area (µm^2^) of spores at time zero that went on to successfully undergo the first cell division. The scripts we generated are all publicly available in Stowers ODR at https://www.stowers.org/research/publications/LIBPB-2559.

### Electron Microscopy

We stored spores at 25 or 42°C for varying amounts of time and then completed either Scanning Transmission Electron Microscopy (STEM) or Scanning Electron Microscopy (SEM). For STEM, we fixed spores in 1.5% KMnO_4_ on ice for 30 minutes, rinsed with ultrapure water five times for 10 minutes each and embedded the spores in 1% low melt agarose (*Frankl et al., 2015*). We cut the agarose block into pieces less than 1mm^3^, prior to serial ethanol dehydration. The dehydration steps were for 15 minutes at room temperature for 30, 50, 70, 80, 90 and three 100% incubations. We then moved the samples into Hard Plus resin (Electron Microscopy Sciences) with accelerator starting with a 25% incubation for 2.5 hours then into a 50% incubation overnight at room temperature, 75% the next day, then 100% and microwaved (Pelco) for 250W for 3 minutes and incubated over the next night. The next day, we did 3 changes of 100% resin, all microwaved at the same settings before embedding and hardening at 60°C for 48 hours. We cut sections at 80nm using a Leica Actos ultramicrotome and post-stained the grids 3 minutes with 1% Sato’s Triple Lead, 4 minutes with 4% uranyl acetate in 70% methanol, and 5 minutes with 1% Sato’s Triple Lead. We imaged the grids on a Zeiss Merlin SEM with STEM detector at 30kV 700pA.

For SEM, we fixed spores in 50mM fixative containing 2.5% glutaraldehyde, 2% paraformaldehyde, 1mM CaCl2, 1% sucrose buffered with 50mM Na Cacodylate with pH 7.4 at 25°C for two hours and stored the spores in fixative overnight at 4°C. We then processed the spores with a TOTO protocol (*Jongebloed et al., 1999*). Briefly, we rinsed the spores five times for 5 minutes each in ultrapure water, incubated in 1% Tannic Acid for 1 hour at room temperature on a nutator, washed with the same regime as above, incubated in 1% aqueous osmium tetraoxide for 1 hour at room temperature on a nutator, washed again as above, incubated in 2% filtered aqueous thiocarbohydrizide for an hour at room temperature on a nutator, washed again, followed by another hour in 1% osmium tetraoxide for an hour at room temperature. After a final rinse, we dehydrated the spores in ethanol for 10 min through 30, 50, 70, 90 and three 100% washes and critical point dried with a Tousimis Samdri-795. We then mounted a section of kim wipe on a stub with carbon tape and sputter coated (Leica ACE 6000) with 4nm of Au/Pd. The spores were then sprinkled onto the stub which was further coated with 7nm of carbon to mitigate spore charging with little to no additional surface texture. We then imaged the spores at 3kV, 50pA with an In Lens SE2 detector on a Zeiss Merlin SEM and processed the images using Fiji (*Schindelin et al., 2012*) and the CLAHE function (*Zuiderveld, 1994*) with the following parameters: 127, 256, 1.5.

### Stress tolerance experiments

We stored spores at 25, 37, or 42°C for varying amounts of time and then completed the following stress experiments. For heat stress, we plated dilutions of spores on YEAS plates before heat stress and after being placed at 55°C for 1 or 2 hours. We then grew the plates at 32°C for 3 days and counted the number of CFUs and counted again at 5 days to catch any slow-growing colonies. We divided the number of CFUs on the heat-shocked plate by the number of CFUs on the untreated, control plate to calculate the percent survival and averaged the three replicates per each sample. For UV stress, we plated dilutions of spores onto 2 YEAS plates. One of plates was not subjected to UV (control), but the other we exposed to 2000 or 2500 Jules/m^2^ of UV light (254 nm). We then grew the plates at 32°C for 3 days, counted the number of CFUs, and counted again at 5 days. We divided the number of CFUs on the UV-treated plate by the number of CFUs on the untreated, control plate to calculate the percent survival.

## Data availability

All the reagents used in this study are available upon request. Original data underlying this manuscript can be accessed from the Stowers Original Data Repository at https://www.stowers.org/research/publications/LIBPB-2559. This includes all raw images, as well as the Deep Learning scripts used for image analysis and plotting.

## Acknowledgments

This work was funded by the Stowers Institute for Medical Research (SEZ) and NIH NIGMS grants DP2 GM132936 and R35 GM151982 (SEZ). The funders have no role in study design, data collection and analysis, decision to publish, or preparation of the manuscript. All authors received salary support from the Stowers Institute for Medical Research. The authors disclose no competing interests.

---

**Figure S1.**
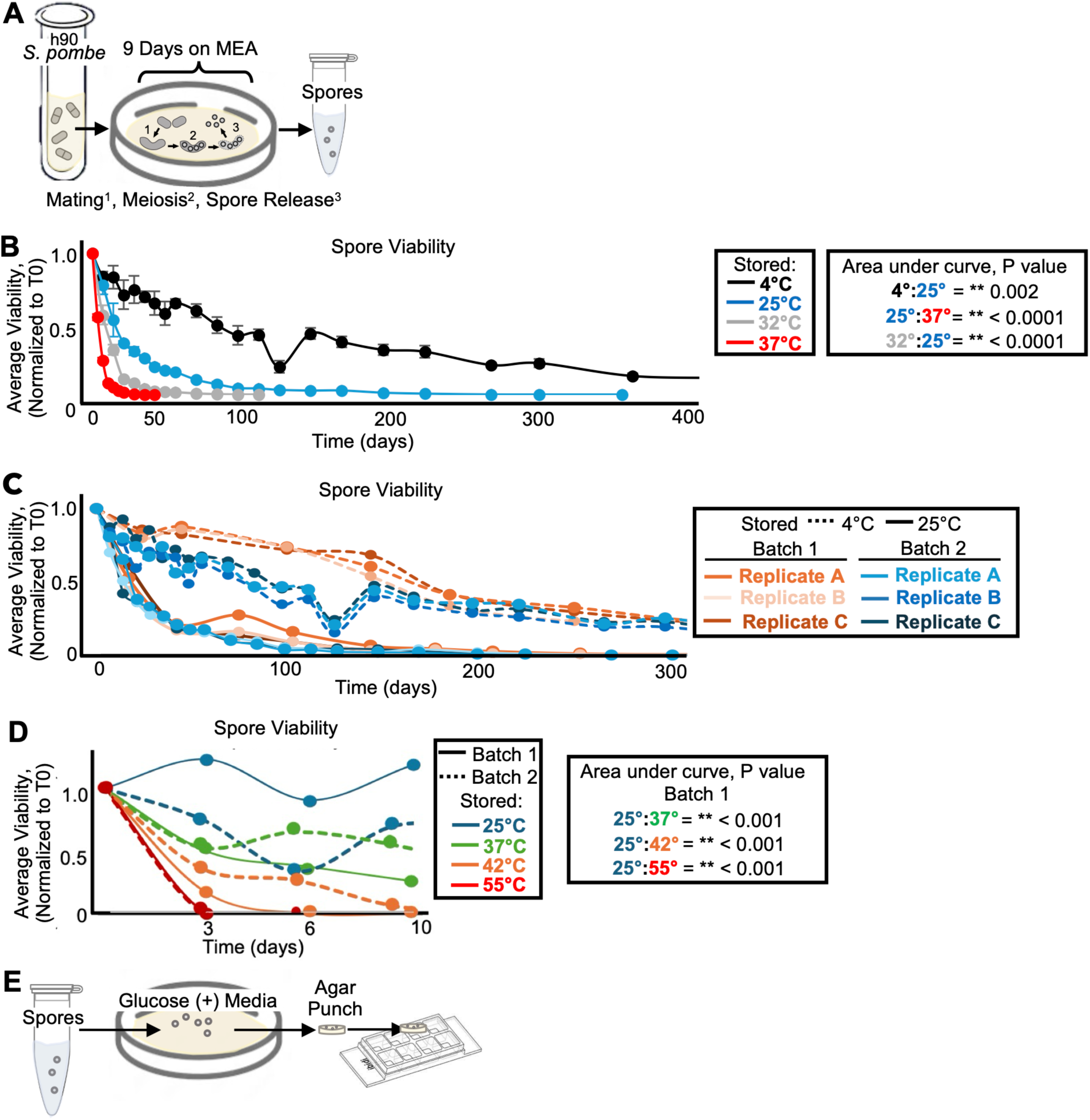
Storage temperature affects spore health and longevity. (A) Cartoon of the spore isolation approach used for all experiments (see methods for more details). A haploid strain proficient in mating-type switching (h90) was cultured in rich media (YES) then plated on sporulation media (MEA) at 25°C for 9 days. The spores were then scraped into tubes and the remaining vegetative cells were killed with glusulase and ethanol treatment. (B) One population of spores stored in water was split and stored at 4°C, 25°C, 32°C, or 37°C. The stored spores were then sampled over time to assay the number of colony-forming units (CFUs) on rich media (YEAS), which were normalized to the time zero starting point. The average of 3 replicates is shown and the error bars represent standard deviations. ** indicates p value < 0.01, comparing areas under the curves via t-test. These are the same data as shown in Figure 1A with more timepoints displayed. (C) This plot follows the same format as (B), but shows the variability present within and between experiments. Batch 1 (orange shades) is a distinct experiment from that shown in (B) and each of three replicates of spores stored at 4°C (dotted lines) and 25°C (solid lines) is shown. Batch 2 (blue shades) is the same data as shown in (B) for spores stored at 4°C (dotted lines) and 25°C (solid lines), except each individual replicate is plotted. (D) An independent, but analogous experiment as that presented in (B) and (C) where the viability of spores stored at 25°C, 37°C, 42°C, and 55°C was assayed over time in two batches. (E) To prepare cells for imaging via timelapse microscopy experiments in this study, we placed spores on agarose plates (YEAS or PMG). We then took a punch of this agar plate and transferred the agar plug to an 8-well Ibidi dish, placing it upside down, trapping the spores between the plug and dish.

**Figure S2.**
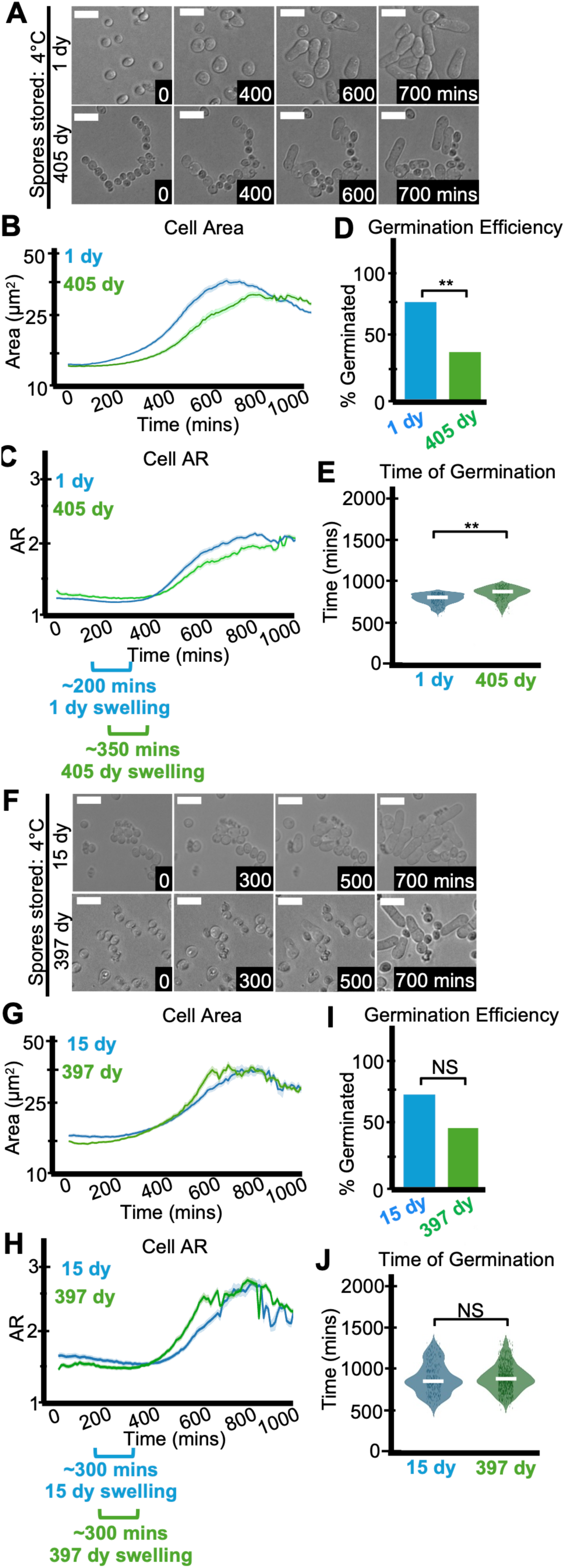
Storage temperature affects spore health. (A-E) Imaging analyses of spores stored at 4°C for one day or 405 days before being plated on rich media (YEAS) to induce germination at 32°C. Cells were imaged every 10 minutes for 1000 minutes. (A) Timelapse microscopy images of germinating spores. The time in minutes is displayed and the scale bars represent 10 μm. (B) Cell area (µm^2^) and (C) cell aspect ratio (AR) during germination. The shaded area around each line represents standard error. In some instances, the standard error is thinner than the line. Note that these cell areas provided by the deep learning algorithm are slightly larger than the actual cell areas (see methods). (D) The fraction of spores manually observed to germinate by the end of the 1,000-minute timelapse. N>50 spores per sample. ** indicates p-value < 0.01 via Fisher’s exact test. (E) Time of first division (when the AR of a cell exceeded 3). ** indicates p-value <0.01 via ANOVA test. N>100 spores per sample. The experiments in (F-J) duplicate those shown in (A-E), but the spores were stored for 15 days or 397 days.

**Figure S3.**
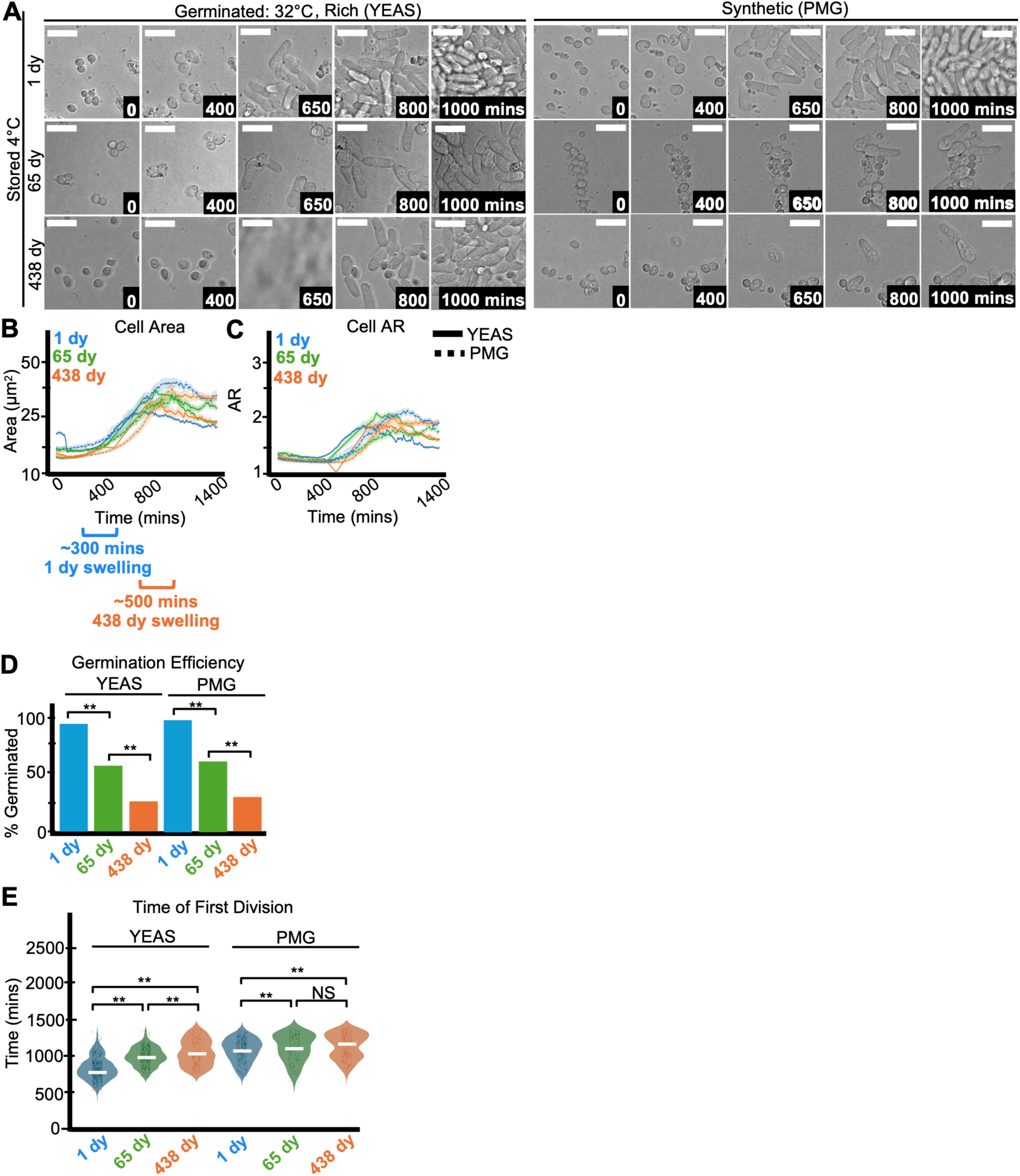
Storage temperature affects spore health. Imaging analyses of spores stored at 4°C for one, 65, or 438 days before being plated on media to induce germination at 32°C. Cells were imaged every 10 minutes for 1,400 minutes. (A) Timelapse microscopy images of spores germinating on rich media (YEAS, left) and synthetic media (PMG, right). The time in minutes is displayed and the scale bars represent 10 μm. The 438-day-old sample on YEAS lost focus at the 650-minute time frame (see methods for details on how such samples were scored). (B) Cell area (µm^2^) and (C) cell aspect ratio (AR) during germination on rich media (YEAS; solid lines) and synthetic media (PMG; dotted lines). The shaded area around each line represents standard error. In some instances, the standard error is thinner than the line. Note that these cell areas provided by the deep learning algorithm are slightly larger than the actual cell areas (see methods). (D) The fraction of spores manually observed to germinate on YEAS and PMG by the end of the 1,400-minute timelapse. N>50 spores per sample. ** indicates p-value < 0.01, Fisher’s exact test. (E) Time of first division (when the AR of a cell exceeded 3) on YEAS and PMG. ** indicates p-value <0.01 via ANOVA test. N>100 spores per sample.

**Figure S4.**
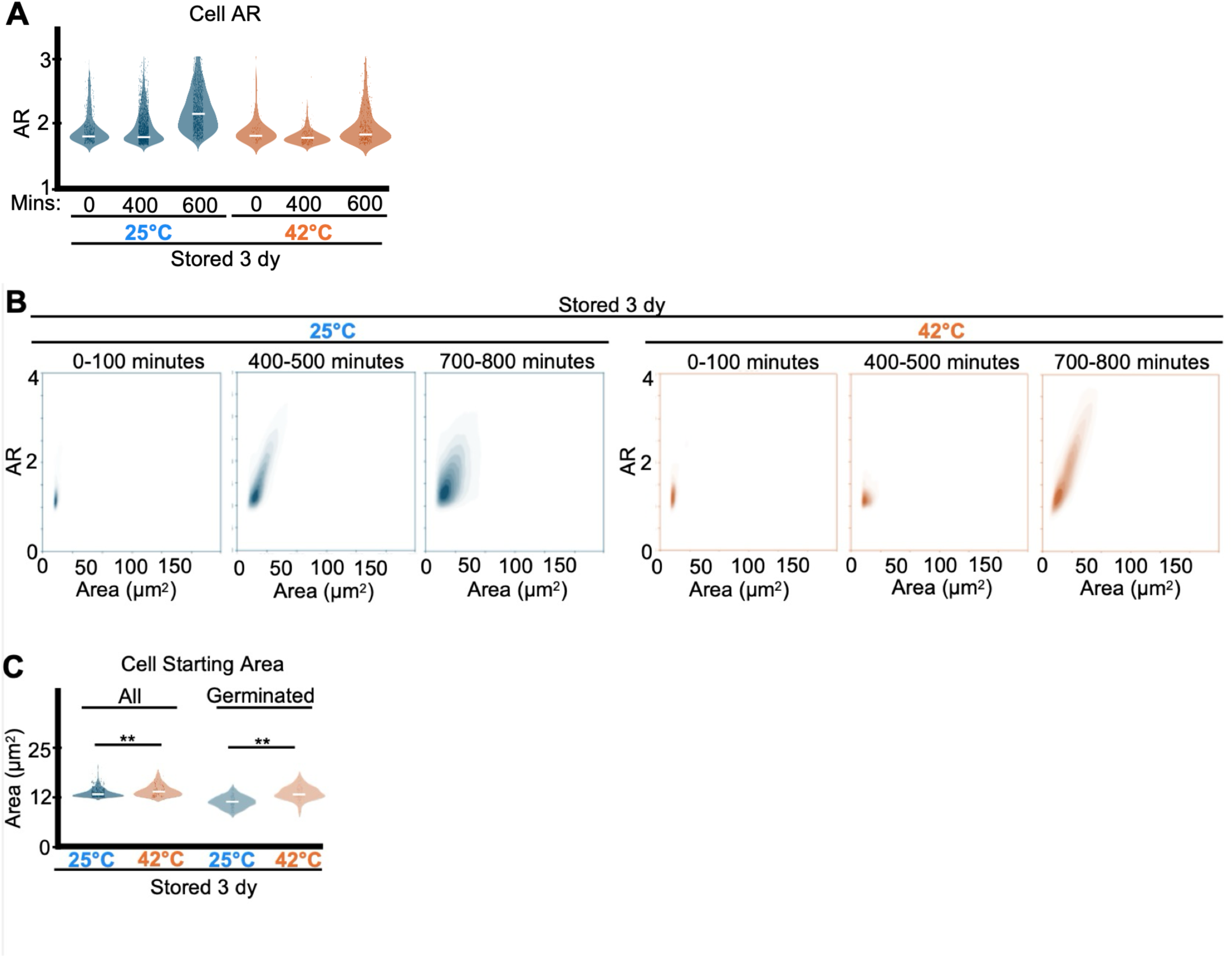
Storage temperature affects spore health. The data in A-B are the same data as those shown in Figure 1C-D but are plotted differently (individual data points rather than population averages) to more clearly illustrate how cell aspect ratio and area change over the course of germination. (A) Aspect Ratio (AR) over time of spores germinating on rich media (YEAS) at 32°C after previous storage of 3 days at 25°C or 42°C in water. (B) Two-dimensional histograms showing the AR and cell area of the entire population of spores analyzed in panel (A). Contour lines were added to help visualize concentrations of cells. (C) The starting area (µm^2^) of all spores (left) or of only the successfully germinated spores analyzed in panel (A). N>50 spores per sample. ** indicates p-value < 0.01 via ANOVA test.

**Figure S5.**
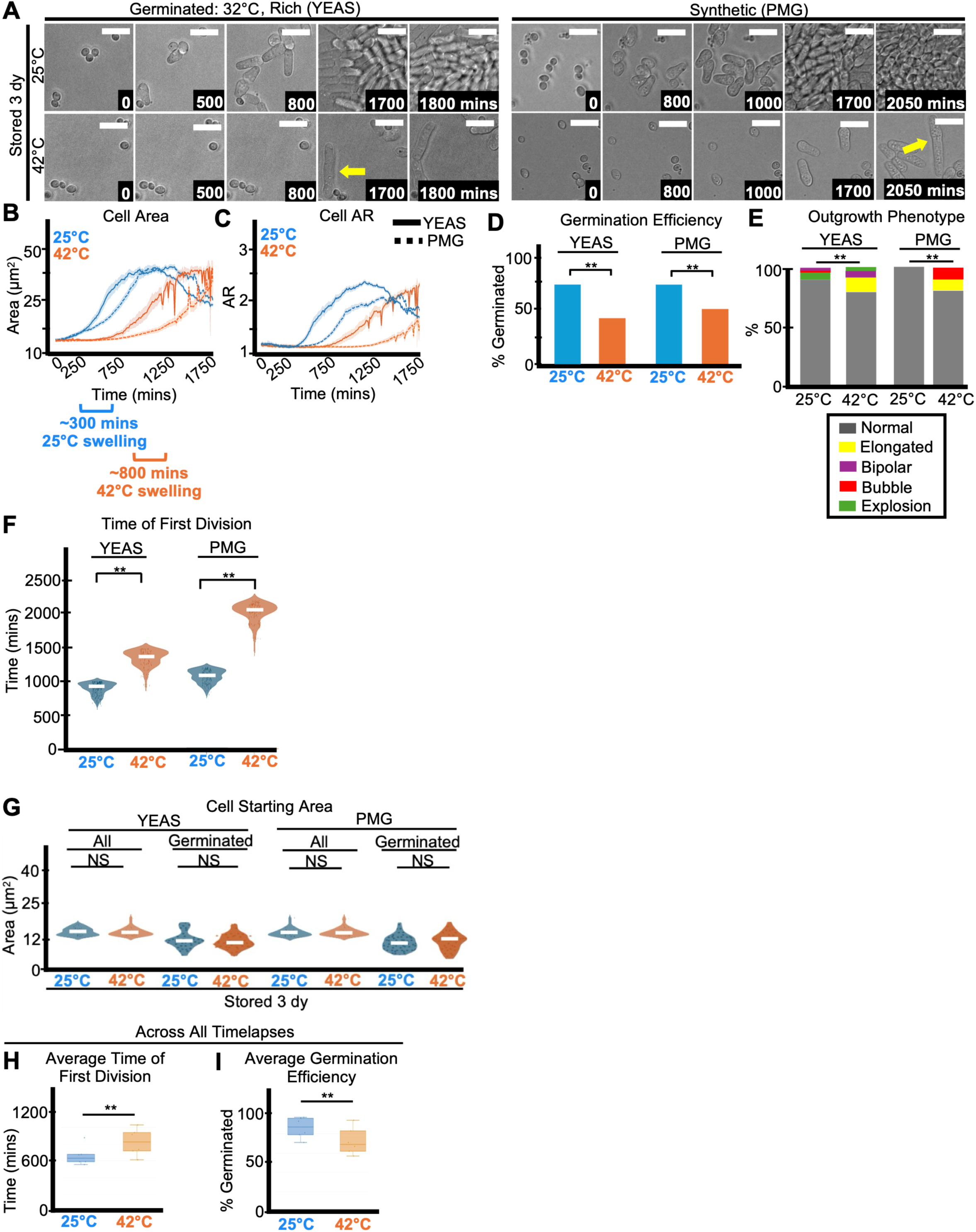
Storage temperature affects spore health. Imaging analyses of spores stored for three days at 25°C or 42°C before being plated on media to induce germination at 32°C. Cells were imaged every 10 minutes for 1,750 minutes. (A) Timelapse microscopy of spores germinating on rich media (YEAS, left) and synthetic media (PMG, right). The yellow arrows illustrated an elongated germination phenotype. The time in minutes is displayed and the scale bars represent 10 μm. (B) Cell area (µm^2^) and (C) cell aspect ratio (AR) during germination on rich media (YEAS; solid lines) and synthetic media (PMG; dotted lines). The shaded area around each line represents standard error. In some instances, the standard error is thinner than the line. Note that these cell areas provided by the deep learning algorithm are slightly larger than the actual cell areas (see methods). (D) The fraction of spores manually observed to germinate on YEAS and PMG by the end of the 1,750-minute timelapse. N>50 spores per sample. ** indicates p-value < 0.01 via Fisher’s exact test. (E) The percentage of spores displaying the indicated outgrowth phenotypes on YEAS and PMG. N>50 outgrowths per sample. ** indicates p-value < 0.01 via Fisher’s exact test. (F) Time of first division (when the AR of a cell exceeded 3) on YEAS and PMG. ** indicates p-value <0.01 via ANOVA test. N>100 spores per sample. (G) Starting area (µm^2^) of all (left) or of only the successfully germinated (right) spores on YEAS and PMG. N>50 spores per sample. ** indicates p-value < 0.01 via ANOVA test. (H) Average time of first division and (I) Average germination efficiency of spores stored for 2-4 days at 25°C (N=6) or 42°C (N=6) across all timelapses with various germination temperatures and media.

**Figure S6.**
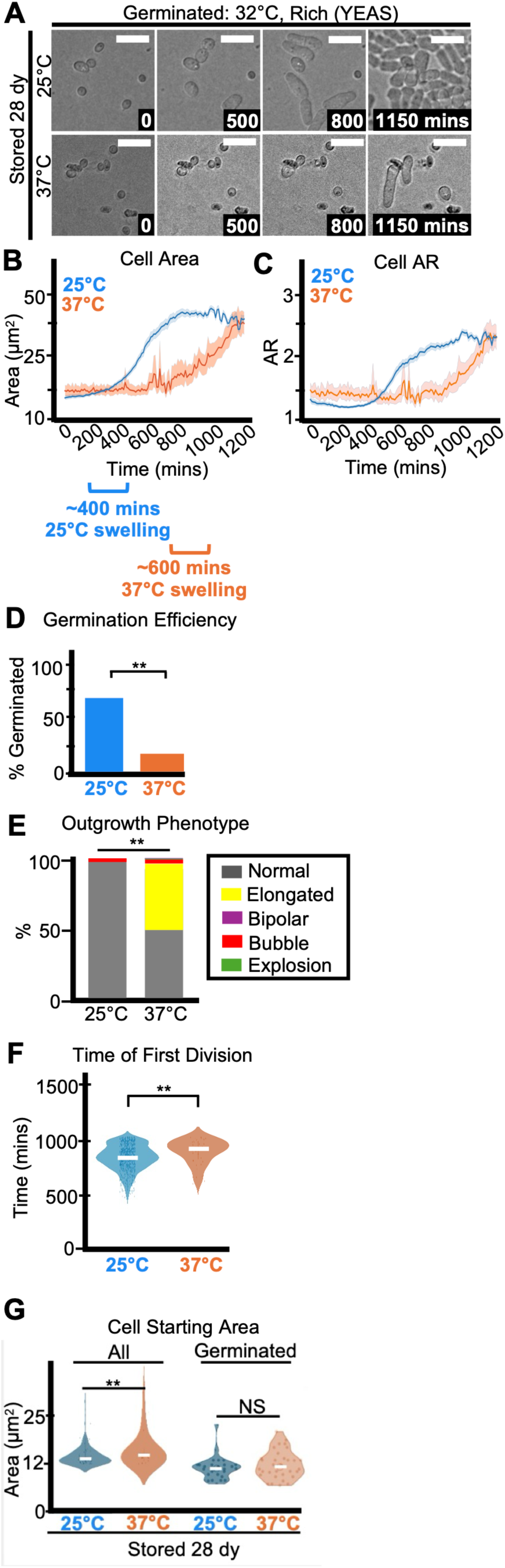
Storage temperature affects spore health. Imaging analyses of spores stored for 28 days at 25°C or 37°C before germination was induced at 32°C on rich media (YEAS). Cells were imaged every 10 minutes for 1,200 minutes. (A) Timelapse microscopy images of germinating spores. The time in minutes is displayed and the scale bars represent 10 μm. (B) Cell area (µm^2^) and (C) cell aspect ratio (AR) during germination. The shaded area around each line represents standard error. In some instances, the standard error is thinner than the line. Note that these cell areas provided by the deep learning algorithm are slightly larger than the actual cell areas (see methods). (D) The fraction of spores manually observed to germinate by the end of the 1,200-minute timelapse. N>50 spores per sample. ** indicates p-value < 0.01, Fisher’s exact test. (E) The percentage of spores displaying the indicated outgrowth phenotypes. N>50 outgrowths per sample. ** indicates p-value < 0.01, Fisher’s exact test. (F) Time of first division (when the AR of a cell exceeded 3). ** indicates p-value <0.01 via ANOVA test. N>100 spores per sample. (G) Starting area (µm^2^) of all (left) or of only successfully germinated (right) spores. N>50 spores per sample. ** indicates p-value < 0.01 via ANOVA test.

**Figure S7.**
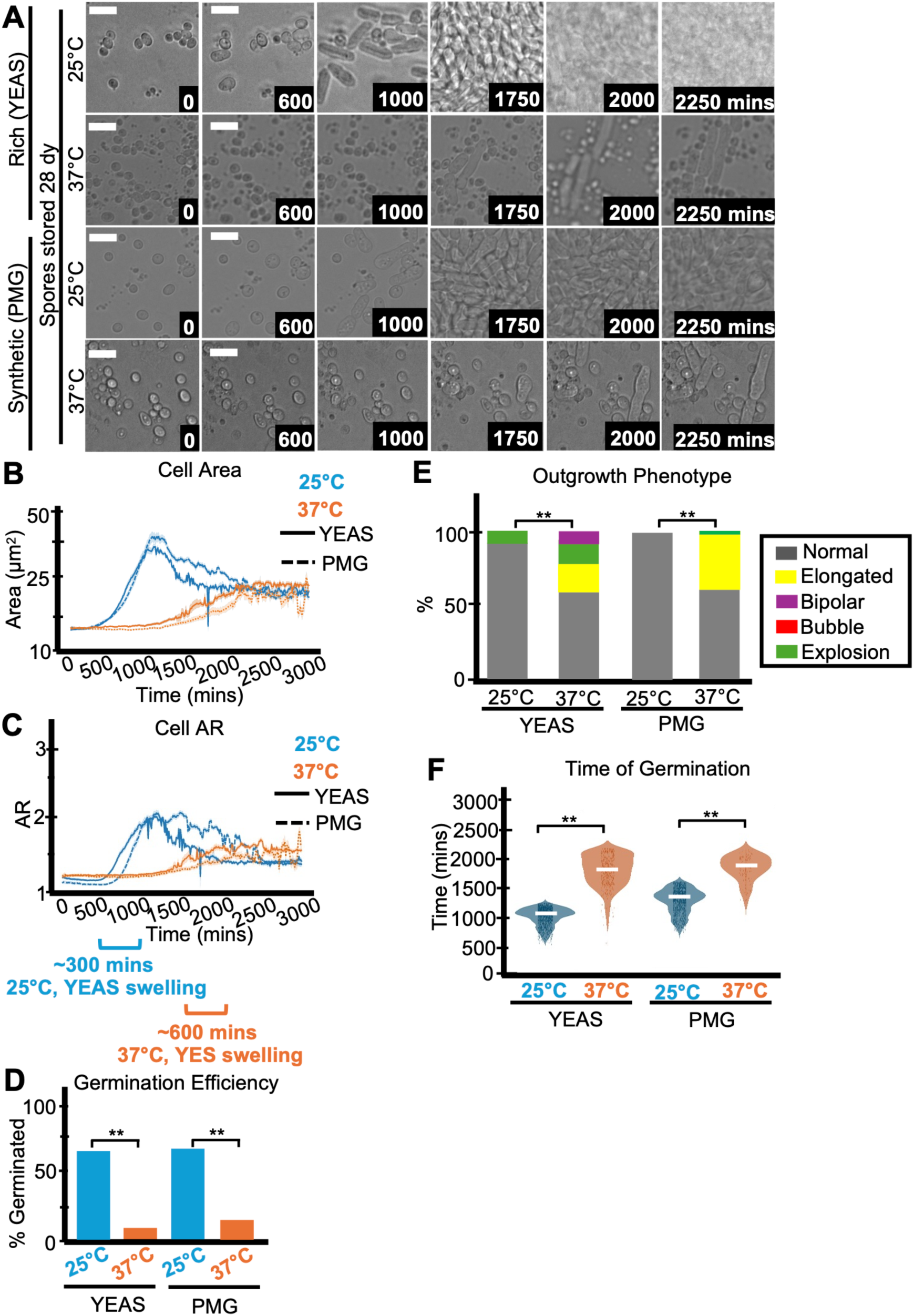
Storage temperature affects spore health. Imaging analyses of spores stored 28 days at 25°C or 37°C before germination was induced on rich media (YEAS) or synthetic media (PMG) at 32°C. The cells were imaged every 10 minutes for 3,000 minutes. (A) Timelapse microscopy images of cells during germination. The storage temperature and germination medium are listed to the right of each row of images. The time in minutes is displayed and the scale bars represent 10 μm. (B) Cell area (µm^2^) and (C) cell aspect ratio (AR) during germination. The shaded area around each line represents standard error. In some instances, the standard error is thinner than the line. Note that these cell areas provided by the deep learning algorithm are slightly larger than the actual cell areas (see methods). (D) The fraction of spores manually observed to germinate by the end of the 3000-minute timelapse. N>50 spores per sample. ** indicates p-value < 0.01 via Fisher’s exact test. (F) Time of first division (when the AR of a cell exceeded 3). ** indicates p-value <0.01 via ANOVA test. N>100 spores per sample. (E) The percentage of spores displaying the indicated outgrowth phenotypes. N>50 outgrowths per sample. ** indicates p-value < 0.01 via Fisher’s exact test.

**Figure S8.**
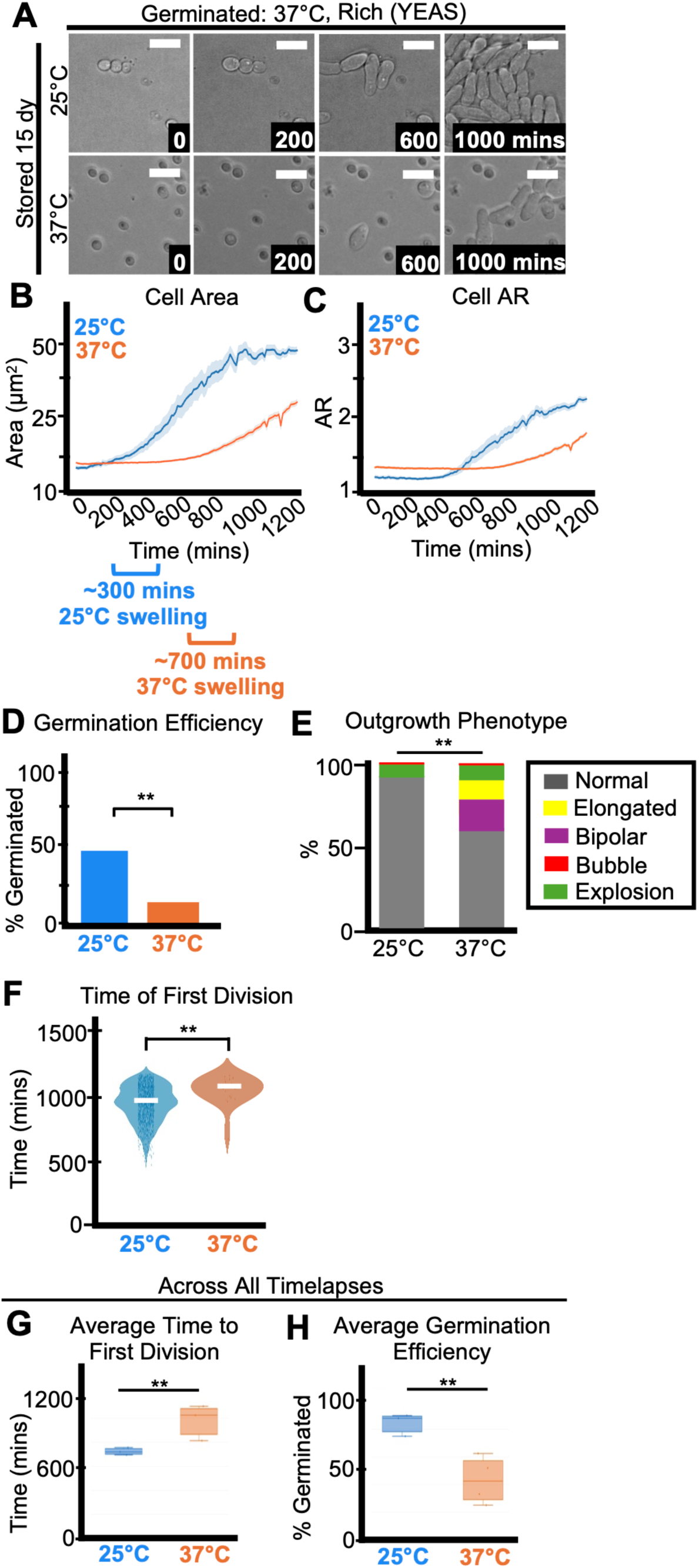
Storage temperature affects spore health. Imaging analyses of spores stored for 15 days at 25°C or 37°C before germination was induced at 37°C on rich media (YEAS). Cells were imaged every 10 minutes for 1,200 minutes. (A) Timelapse microscopy images of germinating spores. The time in minutes is displayed and the scale bars represent 10 μm. (B) Cell area (µm^2^) and (C) cell aspect ratio (AR) during germination. The shaded area around each line represents standard error. In some instances, the standard error is thinner than the line. Note that these cell areas provided by the deep learning algorithm are slightly larger than the actual cell areas (see methods). (D) The fraction of spores manually observed to germinate by the end of the 1,200-minute timelapse. N>50 spores per sample. ** indicates p-value < 0.01 via Fisher’s exact test. (E) The percentage of spores displaying the indicated outgrowth phenotypes. N>50 outgrowths per sample. ** indicates p-value < 0.01 via Fisher’s exact test. (F) Time of first division (when the AR of a cell exceeded 3). ** indicates p-value <0.01 via ANOVA test. N>100 spores per sample. (G) Average time of first division and (I) Average germination efficiency of spores stored for 15-35 days at 25°C (N=9) or 37°C (N=4) across all timelapses with various germination temperatures and media.

**Figure S9.**
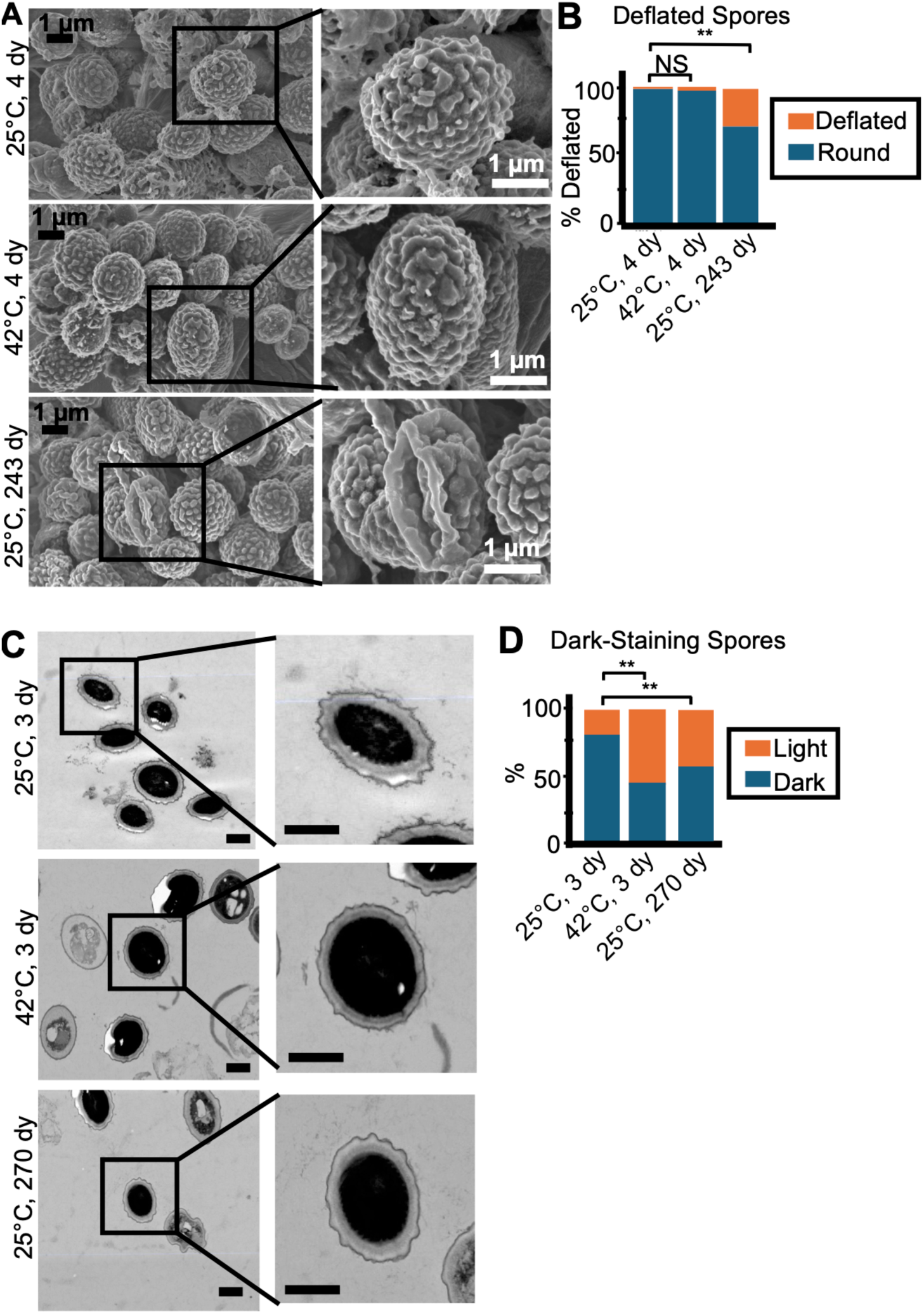
Storage temperature affects spore health. (A) Scanning electron microscopy images of spores stored for 4 days at 25°C (top), 4 days at 42°C (middle), and 243 days at 25°C (bottom). The inset on the 243-day old sample centers on a ‘deflated’ spore. Scale bars represent 1 μm. (B) Percentage of deflated and round spores observed in samples represented in panel (A). N> 50 spores. ** indicates p-value < 0.01 via Fisher’s exact test. (C) Scanning transmission electron microscopy images of spores stored for 3 days at 25°C (top), 3 days at 42°C (middle), or 270 days at 25°C (bottom). Each of the insets focus on a ‘dark-staining’ spore, but a ‘light-staining’ spore is just to the left of the inset spore in the 42°C sample. Scale bars represent 1 μm. (D) Percentage of spores stored for 3 days at 25°C or 42°C, or 270 days at 25°C with light-staining or dark-staining cytoplasm. N> 50 spores. ** indicates p-value < 0.01 via Fisher’s exact test.

**Figure S10.**
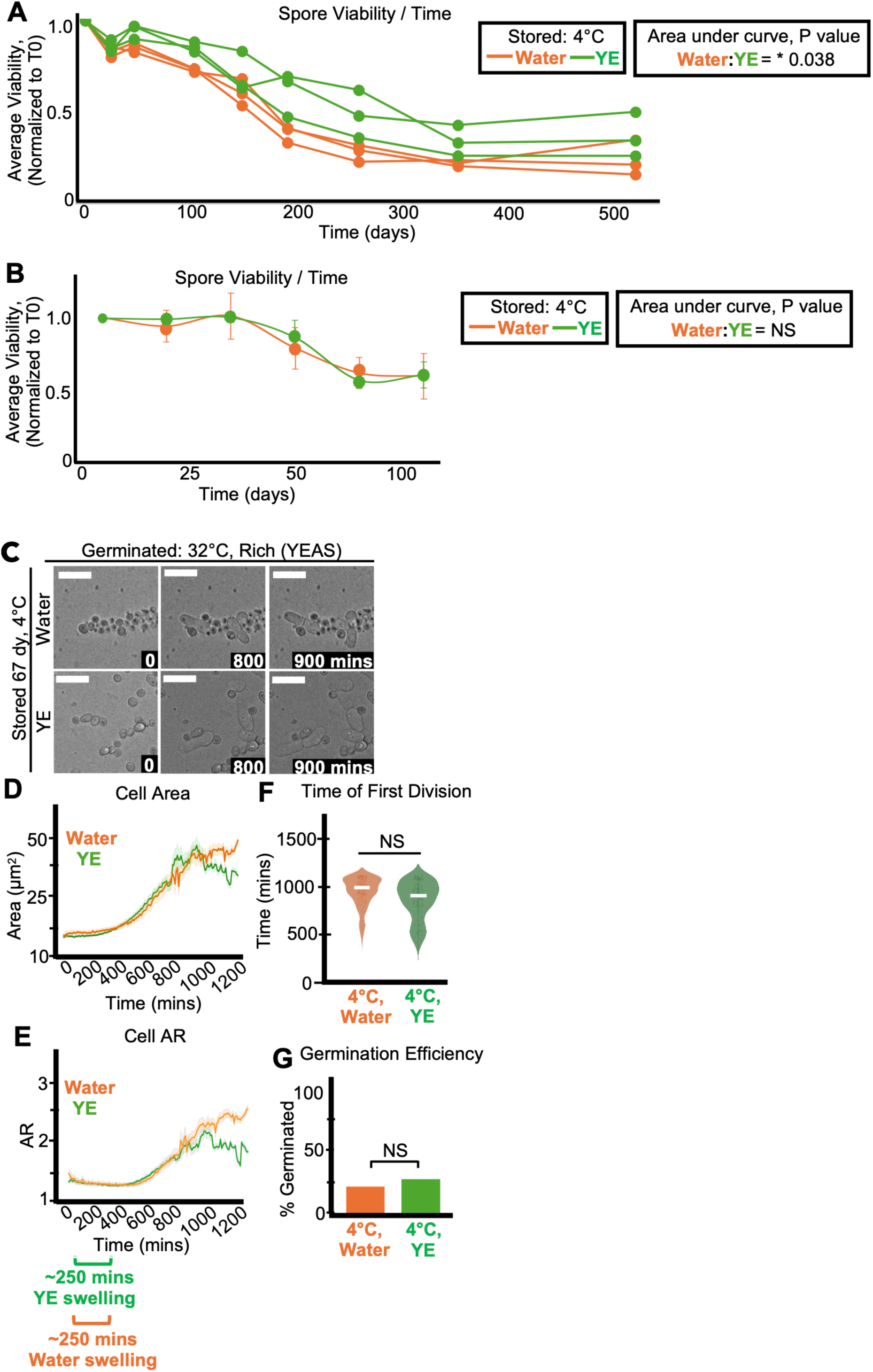
Storage media significantly affects spore longevity, but not health. (A) One population of spores was split and resuspended in either 0.5% yeast extract solution without glucose (YE) or water and then stored at 4°C. Panel (B) shows a repeat of the experiment shown in (A). In both experiments, the stored spores were then sampled over time to assay the number of colony-forming units (CFUs) on rich media (YEAS), which were normalized to the time zero starting point. In (A), three replicates are shown for each storage medium. In (B), The average of 3 replicates is shown and the error bars represent standard deviations. ** indicates p value < 0.01, * indicates p value < 0.05, comparing areas under the curves via t-test. (C-G) Imaging analyses of spores stored for 67 days at 4°C in either water or yeast extract solution before being plated on rich media to induce germination at 32°C. Cells were imaged every 10 minutes for 1,200 minutes. (C) Timelapse microscopy images of germinating spores. The time in minutes is displayed and the scale bars represent 10 μm. (D) Cell area (µm^2^) and (E) cell aspect ratio (AR) during germination. The shaded area around each line represents standard error. In some instances, the standard error is thinner than the line. Note that these cell areas provided by the deep learning algorithm are slightly larger than the actual cell areas (see methods). (F) Time of first division (time when the AR of a cell exceeded 3). NS indicates not significant via ANOVA test. N>100 spores per sample. (G) The fraction of spores manually observed to germinate by the end of the 1,200-minute timelapse. N>50 spores per sample. NS indicates not significant via Fisher’s exact test.

**Figure S11.**
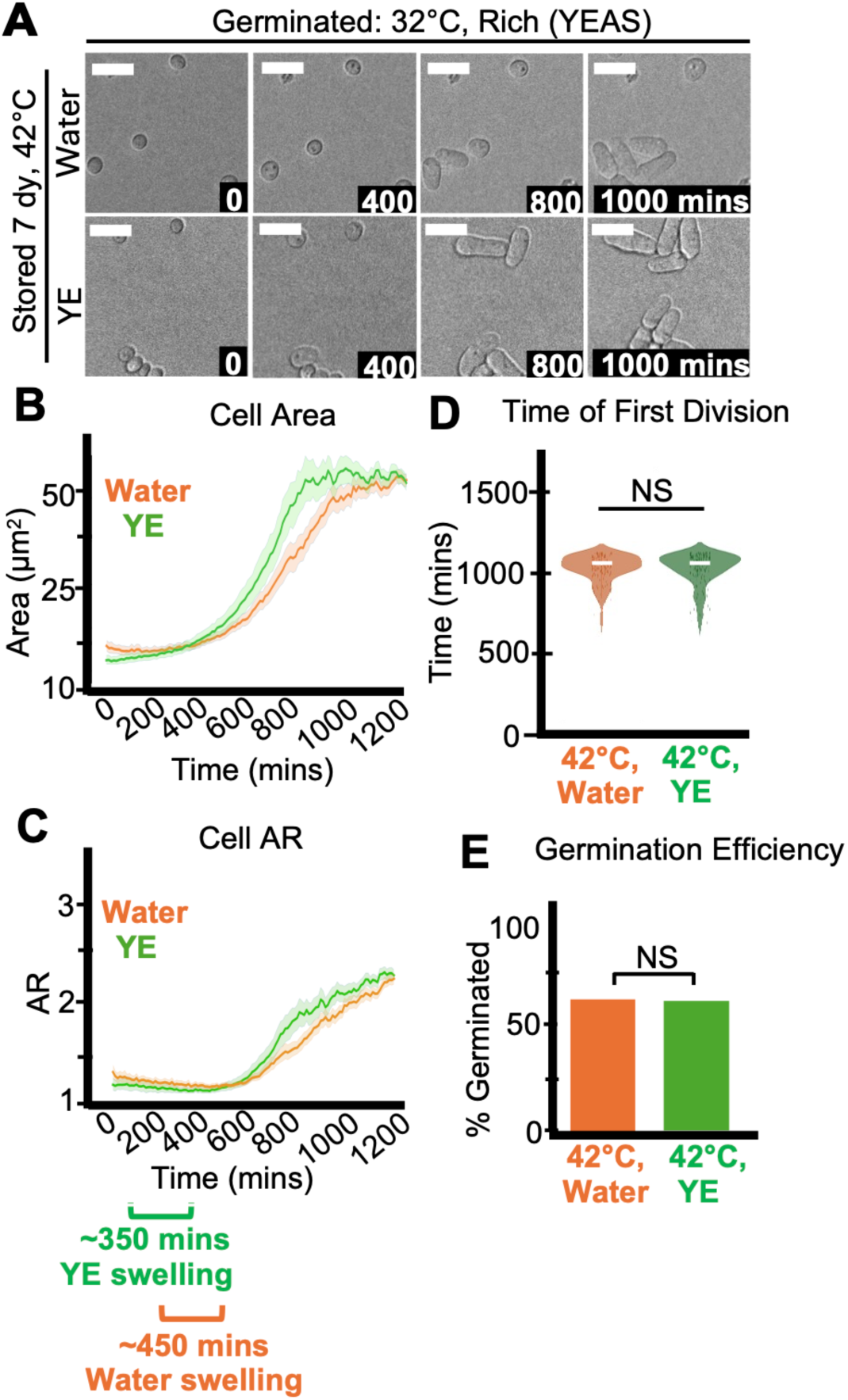
Storage media significantly affects spore longevity, but not health. Imaging analyses of spores stored for 7 days at 42°C in either water or 0.5% yeast extract solution without glucose (YE) before being plated on rich media (YEAS) to induce germination at 32°C. Cells were imaged every 10 minutes for 1,200 minutes. (A) Timelapse microscopy images of germinating spores. The time in minutes is displayed and the scale bars represent 10 μm. (B) Cell area (µm^2^) and (C) cell aspect ratio (AR) during germination. The shaded area around each line represents standard error. In some instances, the standard error is thinner than the line. Note that these cell areas provided by the deep learning algorithm are slightly larger than the actual cell areas (see methods). (D) Time of first division (time when the AR of a cell exceeded 3). NS indicates not significant via ANOVA test. N>100 spores per sample. (E) The fraction of spores manually observed to germinate by the end of the 1,200-minute timelapse. N>50 spores per sample. NS indicates not significant via Fisher’s exact test.

**Figure S12.**
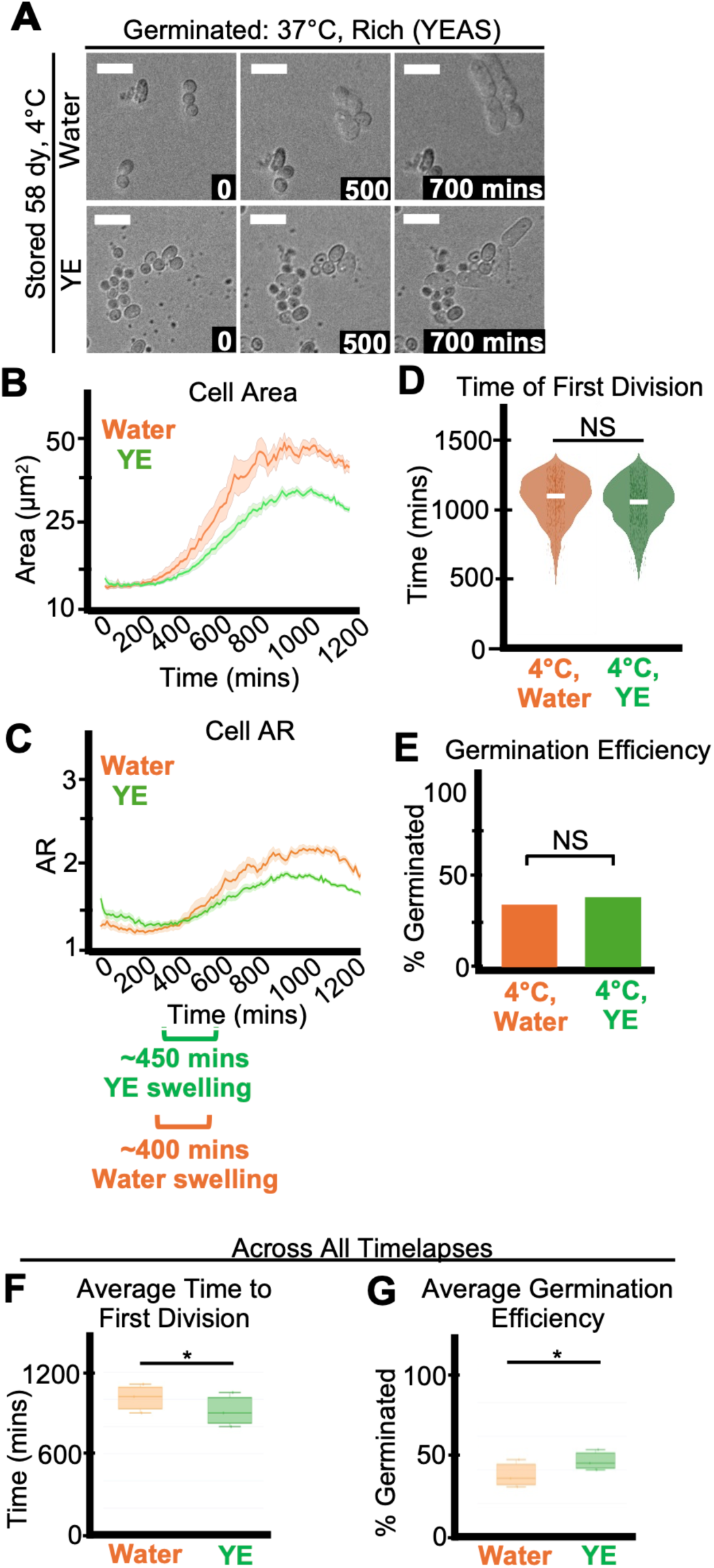
Storage media significantly affects spore longevity, but not health. Imaging analyses of spores stored for 58 days at 4°C in either water or 0.5% yeast extract solution without glucose (YE) before being plated on rich media (YEAS) to induce germination at 37°C. Cells were imaged every 10 minutes for 1,200 minutes. (A) Timelapse microscopy images of germinating spores. The time in minutes is displayed and the scale bars represent 10 μm. (B) Cell area (µm^2^) and (C) cell aspect ratio (AR) during germination. The shaded area around each line represents standard error. In some instances, the standard error is thinner than the line. Note that these cell areas provided by the deep learning algorithm are slightly larger than the actual cell areas (see methods). (D) Time of first division (time when the AR of a cell exceeded 3). NS indicates not significant via ANOVA test. N>100 spores per sample. (E) The fraction of spores manually observed to germinate by the end of the 1,200-minute timelapse. N>50 spores per sample. NS indicates not significant via Fisher’s exact test. (F) Average time of first division and (G) Average germination efficiency of spores stored for 7-67 days in water (orange) or YE (green) at 4°C across all timelapses, including various germination temperatures and media (N=4).

**Figure S13.**
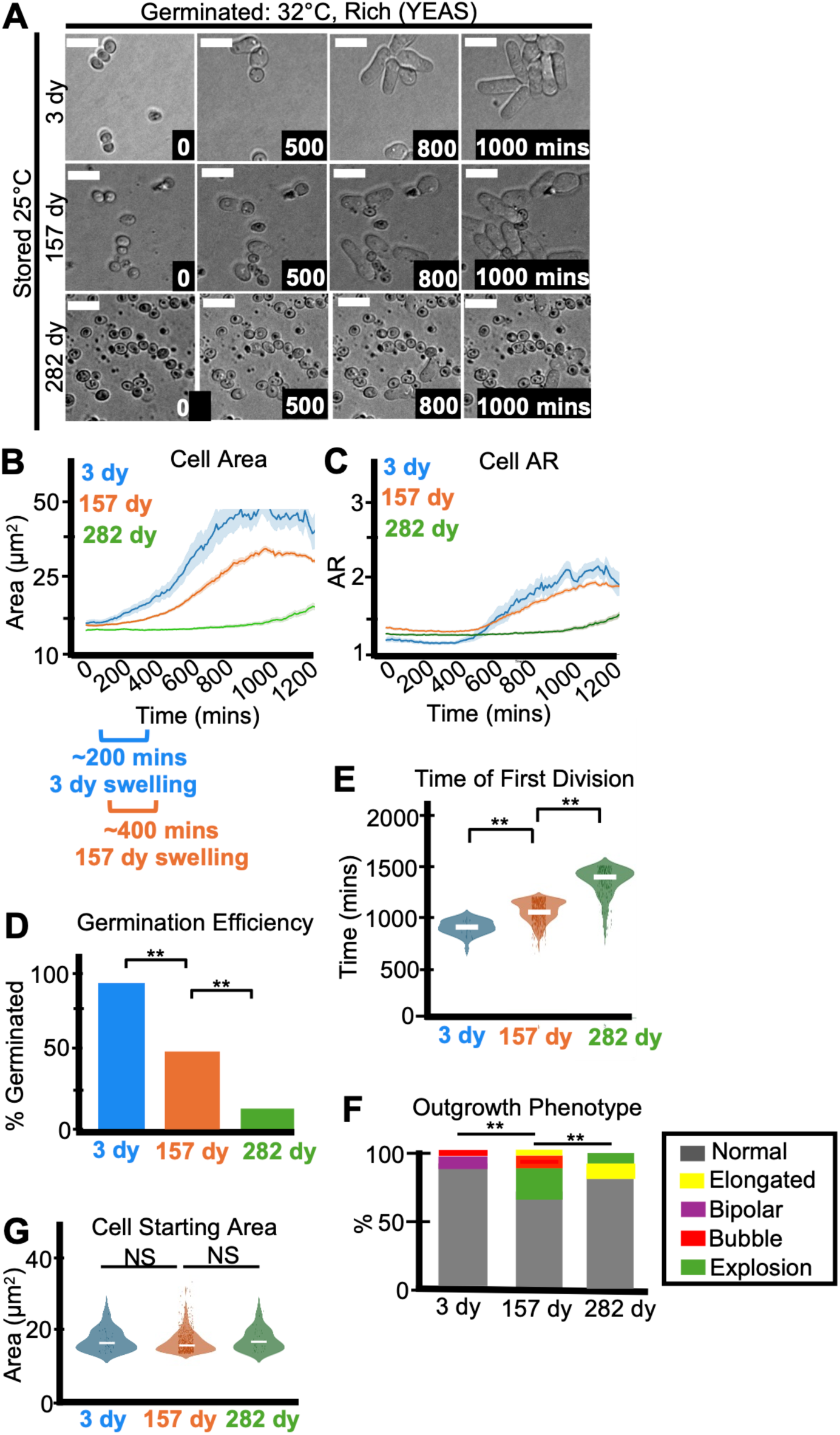
Age affects spore health. (A) Imaging analyses of spores stored for 3, 157, or 282 days at 25°C in water before being plated on rich media (YEAS) to induce germination at 32°C. Cells were imaged every 10 minutes for 1,200 minutes. (A) Timelapse microscopy images of germinating spores. The time in minutes is displayed and the scale bars represent 10 μm. (B) Cell area (µm^2^) and (C) aspect ratio (AR) during germination. The shaded area around each line represents standard error. In some instances, the standard error is thinner than the line. Note that these cell areas provided by the deep learning algorithm are slightly larger than the actual cell areas (see methods). (D) The fraction of spores manually observed to germinate by the end of the 1,200-minute timelapse. N>50 spores per sample. ** indicates p-value < 0.01 via Fisher’s exact test. (E) Time of first division (time when the AR of a cell exceeded 3). ** indicates p-value < 0.01 via ANOVA test. N>100 spores per sample. (F) The percentage of spores displaying the indicated outgrowth phenotypes. N>50 outgrowths per sample. ** indicates p-value < 0.01 via Fisher’s exact test. (G) Starting area (µm^2^) of spores. N>50 spores per sample. NS indicates not significant via ANOVA test.

**Figure S14.**
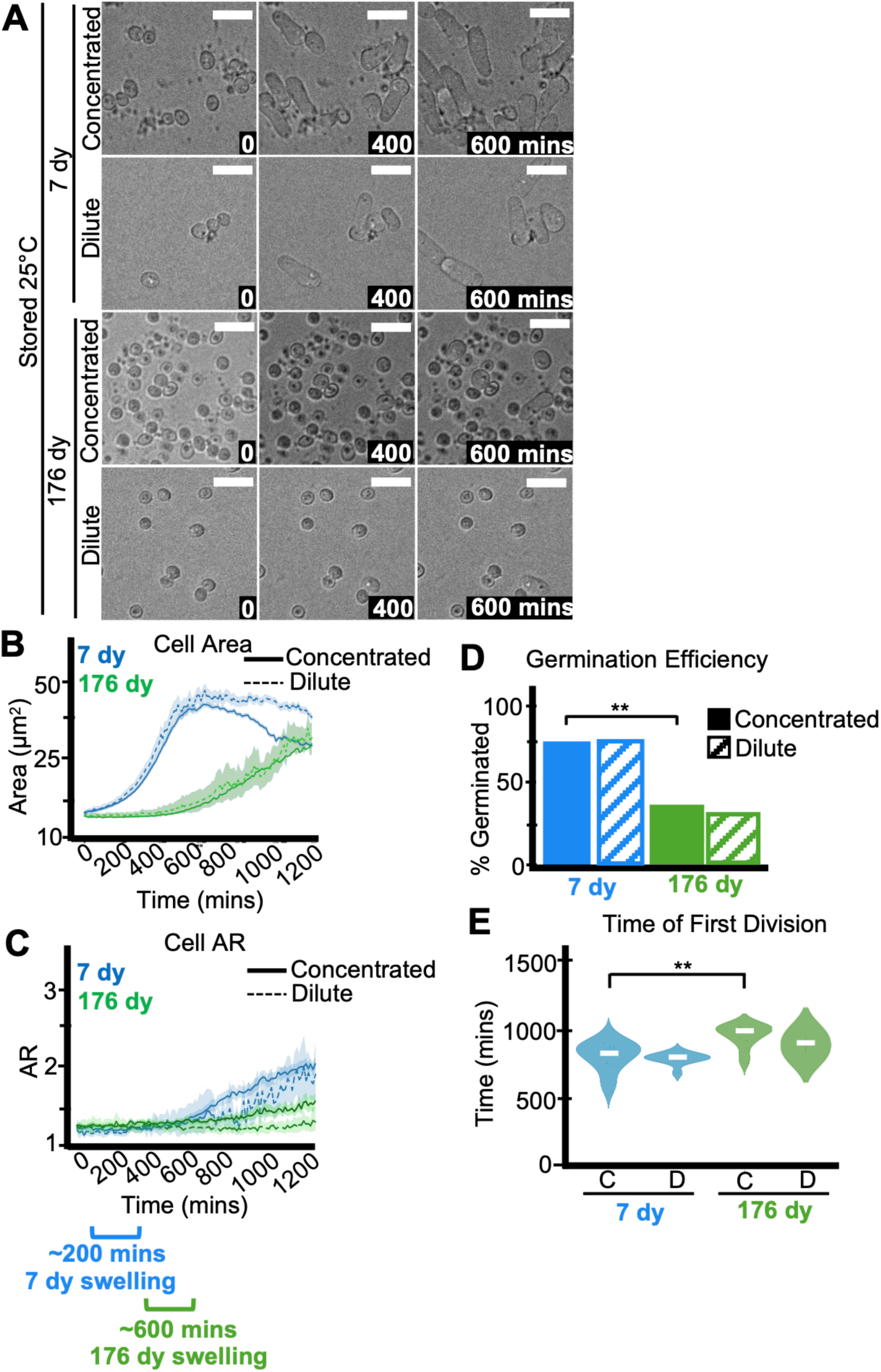
Age affects spore health. Imaging analyses of spores stored for 7 or 176 days at 25°C in water before being plated on rich media (YEAS) in high (concentrated) or low (dilute) concentration to induce germination at 32°C. Cells were imaged every 10 minutes for 1,200 minutes. (A) Timelapse microscopy images of germinating spores. The time in minutes is displayed and the scale bars represent 10 μm. (B) Cell area (µm^2^) and (C) aspect ratio (AR) during germination. The shaded area around each line represents standard error. In some instances, the standard error is thinner than the line. Note that these cell areas provided by the deep learning algorithm are slightly larger than the actual cell areas (see methods). (D) The fraction of spores manually observed to germinate by the end of the 1,200-minute timelapse. N>50 spores per sample. ** indicates p-value < 0.01 via Fisher’s exact test. (E) Time of first division (time when the AR of a cell exceeded 3). ** indicates p-value < 0.01 via ANOVA test. N>100 spores per sample.

**Figure S15.**
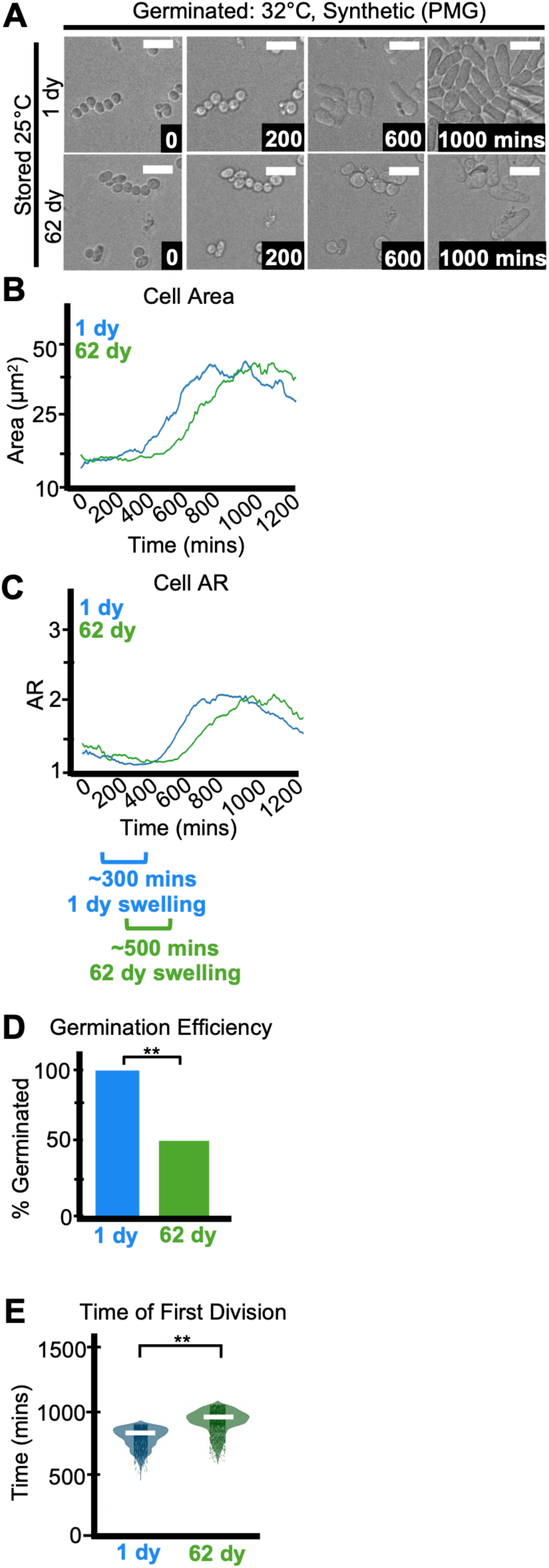
Age affects spore health. Imaging analyses of spores stored for 1 or 62 days at 25°C in water before being plated on synthetic media (PMG) to induce germination at 32°C. Cells were imaged every 10 minutes for 1,200 minutes. (A) Timelapse microscopy images of germinating spores. The time in minutes is displayed and the scale bars represent 10 μm. (B) Cell area (µm^2^) and (C) aspect ratio (AR) during germination. The shaded area around each line represents standard error. In some instances, the standard error is thinner than the line. Note that these cell areas provided by the deep learning algorithm are slightly larger than the actual cell areas (see methods). (D) The fraction of spores manually observed to germinate by the end of the 1,200-minute timelapse. N>50 spores per sample. ** indicates p-value < 0.01 via Fisher’s exact test. (E) Time of first division (time when the AR of a cell exceeded 3). ** indicates p-value < 0.01 via ANOVA test. N>100 spores per sample.

**Figure S16.**
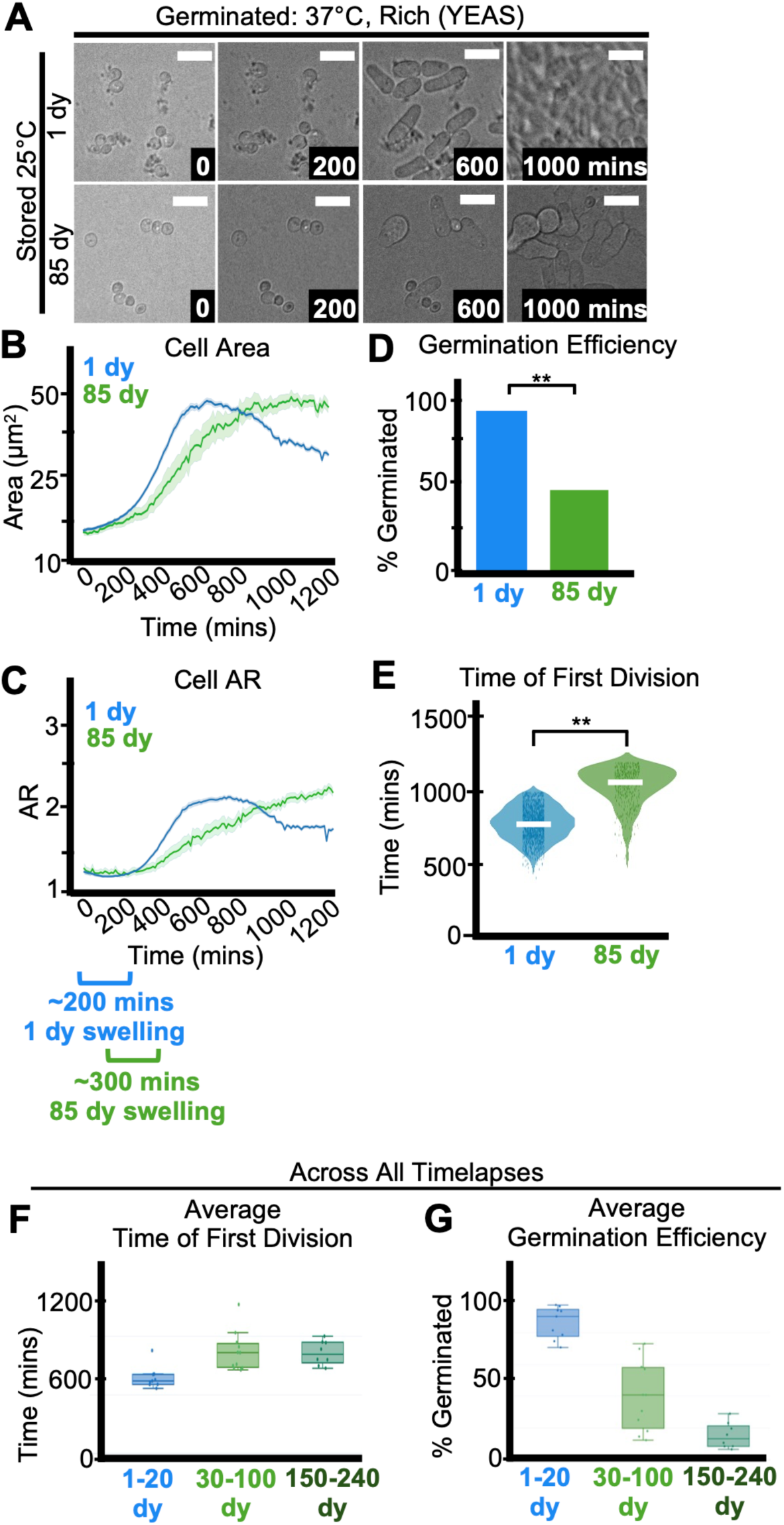
Age affects spore health. Imaging analyses of spores stored for 1 or 85 days at 25°C in water before being plated on rich media (YEAS) to induce germination at 37°C. Cells were imaged every 10 minutes for 1,200 minutes. (A) Timelapse microscopy images of germinating spores. The time in minutes is displayed and the scale bars represent 10 μm. (B) Cell area (µm^2^) and (C) aspect ratio (AR) during germination. The shaded area around each line represents standard error. In some instances, the standard error is thinner than the line. Note that these cell areas provided by the deep learning algorithm are slightly larger than the actual cell areas (see methods). (D) The fraction of spores manually observed to germinate by the end of the 1,200-minute timelapse. N>50 spores per sample. ** indicates p-value < 0.01 via Fisher’s exact test. (E) Time of first division (time when the AR of a cell exceeded 3). ** indicates p-value < 0.01 via ANOVA test. N>100 spores per sample. (F) Average time of first division and (G) Average germination efficiency on YEAS of spores stored for 1-20, 30-100, or 150-240 days in water at 25°C across all timelapses, including various germination temperatures and media (N=4).

**Figure S17.**
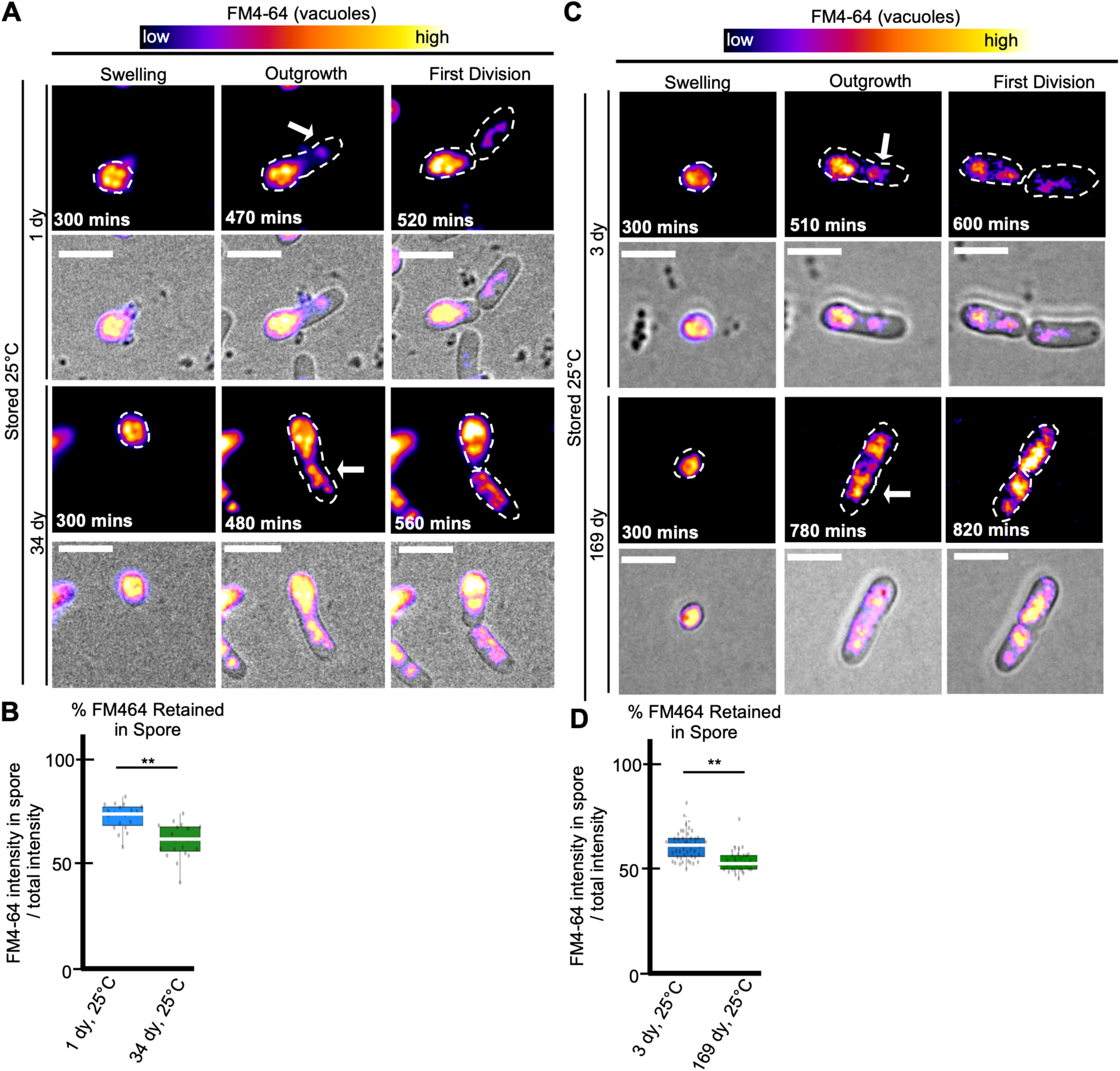
Age affects asymmetry of vacuole segregation during germination. Imaging analyses of vacuole segregation in FM4-64-stained spores previously stored at 25°C in water and then germinated on rich media (YEAS) at 32°C. The cells were imaged every 10 minutes for 20 hours. (A, C) Timelapse microscopy images of FM4-64-stained spores aged for 1 or 34 days (A) and 3 or 169 days (C). The brightness and contrast are not the same for all images but were adjusted so the spore bodies appeared to have similar levels of signal. Scale bars represent 10 μm. (B, D) Percentage of the total FM4-64 signal retained in the spore-body verses the germ tube outgrowth for spores previously aged for 1 or 34 days (B) and 3 or 169 days (D). ** indicates p-value <0.01 via Fisher’s exact test. N> 30 spores.

**Figure S18.**
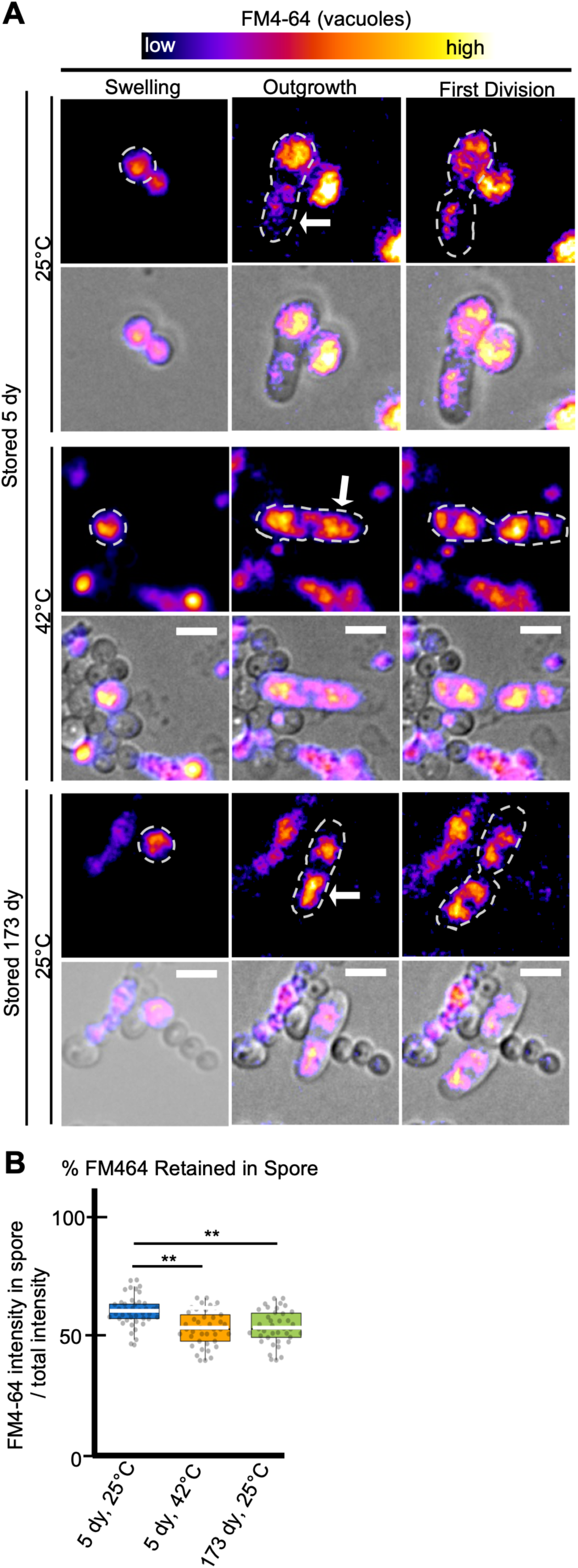
Age affects asymmetry of vacuole segregation during germination. Imaging analyses of vacuole segregation in FM4-64-stained spores previously stored in water for 5 or 173 days at 25°C, or for 5 days at 42°C. (A) Timelapse microscopy images of FM4-64-stained spores germinating on rich media (YEAS) at 32°C. The brightness and contrast are not the same for all images but were adjusted so the spore bodies appeared to have similar levels of signal. Scale bars represent 10 μm. (B) Percentage of the total FM4-64 signal retained in the spore-body verses the germ tube outgrowth. ** indicates p-value <0.01 via Fisher’s exact test. N> 30 spores.

**Figure S19.**
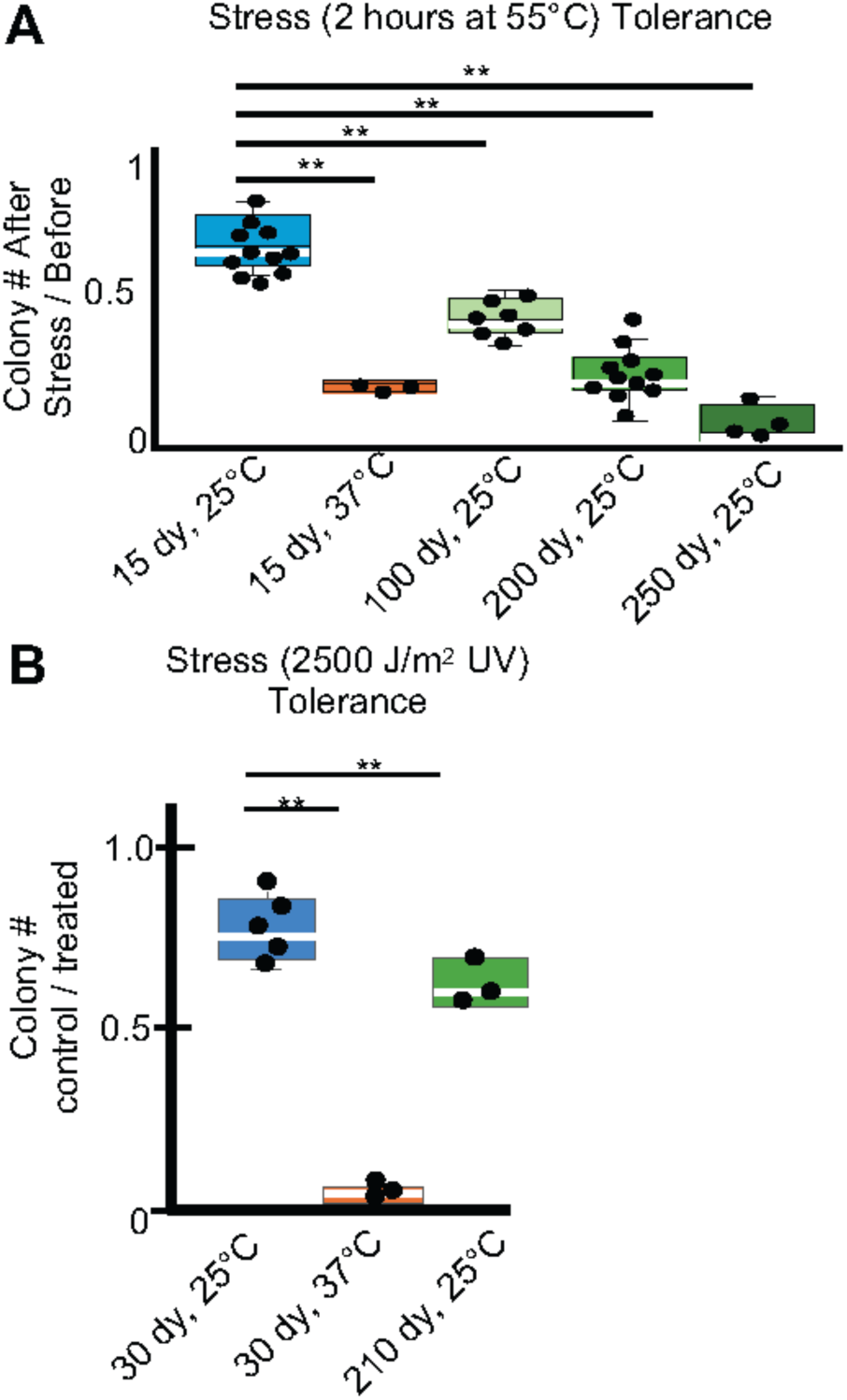
Age and stress affect spore stress tolerance. (A) The colony forming units (CFUs) of spore samples previously stored for 15, 100, 200 or 250 days at 25°C or for 15 days 37°C were assayed by plating on rich media (YEAS) at 32°C both before and after a 2-hour heat shock of 55°C. (B) CFUs of spores samples previously stored for 30 or 210 days at 25°C or 37°C for 30 days before and after exposure to 2500 μJules UV. ** indicates p-value < 0.01 via t-test. N>3 replicates.

